# Efficient modelling of infectious diseases in wildlife: a case study of bovine tuberculosis in wild badgers

**DOI:** 10.1101/2024.01.26.576600

**Authors:** Evandro Konzen, Richard J. Delahay, Dave J. Hodgson, Robbie A. McDonald, Ellen Brooks Pollock, Simon E. F. Spencer, Trevelyan J. McKinley

## Abstract

Bovine tuberculosis (bTB) has significant socio-economic and welfare impacts on the cattle industry worldwide. In the United Kingdom and Ireland, disease control is complicated by the presence of infection in wildlife, principally the European badger. Control strategies tend to be applied to whole populations, but better-targeted management of key sources of transmission, be they individuals or groups, may be more efficient. Mechanistic transmission models can be used to better understand key epidemiological drivers of disease spread and identify high-risk individuals and groups as long as they can be adequately fitted to observed data. However, this is a significant challenge, especially within wildlife populations, because monitoring relies on imperfect diagnostic test information, and even under systematic surveillance efforts (such as capture-mark-recapture sampling) epidemiological events are only partially observed.

To this end we develop a stochastic compartmental model of bTB transmission, and fit this to individual-level data from a unique *>* 40-year longitudinal study of 2,391 badgers using a recently developed individual forward filtering backward sampling algorithm. Modelling challenges are further compounded by spatio-temporal meta-population structures and age-dependent mortality. We develop a novel estimator for the individual effective reproduction number that provides quantitative evidence for the presence of superspreader badgers, despite the population-level effective reproduction number being less than one. We also infer measures of the hidden burden of infection in the host population through time; the relative likelihoods of competing routes of transmission; effective and realised infectious periods; and longitudinal measures of diagnostic test performance. This modelling framework provides an efficient and generalisable way to fit state-space models to individual-level data in wildlife populations, which allows identification of high-risk individuals and exploration of important epidemiological questions about bTB and other wildlife diseases.

**Author summary:** Wild animals commonly harbour infectious diseases with risk of spillover to humans and livestock. We fitted an individual-level stochastic spatial meta-population model of bovine tuberculosis (bTB) transmission to data from a long-term longitudinal study of the European badger (*Meles meles*). Our framework provides an efficient and generalisable means of fitting state-space models to individual-level data, to identify high-risk individuals and explore important epidemiological questions. We develop a novel estimator for the individual effective reproduction number, providing quantitative evidence for the presence of superspreader badgers (those individuals most responsible for onward transmission of infection), despite the population-level effective reproduction number being less than one. Predicting the hidden burden of infection in individuals and social groups is critical for disease management but challenging in practice, since monitoring relies on imperfect surveillance and diagnostic testing, however control of bTB in badgers could be substantially increased by targeting interventions at high-risk groups.

## Introduction

Despite extensive research efforts to evaluate risks of bovine tuberculosis (bTB) transmission between cattle and badgers [1–9], there remains considerable debate about the relative impacts of within- and between-species spread, and the efficacy of control measures such as badger culling. At Woodchester Park (WP) in Gloucestershire, south-west England, a population of wild badgers has been studied since 1978, providing a unique long-term empirical data set that has provided unprecedented insights into badger ecology, bTB infection in badgers, and transmission to cattle [10–14]. These data afford a rare opportunity to help improve our understanding of the dynamics of bTB spread within a wild badger population. To this end we fit a mechanistic compartmental transmission model to individual-level data, capturing the spread and progression of bTB in the badger host population over a 40-year time period. We use the model to infer critical epidemiological parameters including the relative influence of frequency- and density-dependent transmission processes, estimates for the duration of latent and infectious periods, age-dependent mortality risks, and longitudinal changes in diagnostic test performance.

We also have a particular interest in characterising potential heterogeneity among individual badgers in their responsibility for onward transmission-of-infection. Common measures for characterising the propensity of disease spread within a population are the *basic* and *effective* reproduction numbers [15–18]. The population-level reproduction number has been inferred to be close to one in both species [1, 7] implying endemic persistence of infection. However, it has been widely discussed that population-level reproduction numbers ignore the propensity for some individuals to make a disproportionately large contribution to the spread of infection, so-called ‘superspreaders’ [19–25]. Superspreading is rarely characterised in wildlife populations, but can facilitate explosive early spread and transient epidemics, and maintain infection burden even in the absence of widespread transmission. Identifying spatial locations and/or individuals that represent a high transmission risk should inform targeted disease management strategies [26]. For example, if individuals with a high probability of being infected could be removed selectively, then any unintended consequences of population-level disruption (caused for example by non-selective culling, a process which has been linked to enhanced movement and disease spread in culled badger populations [27]) might be ameliorated.

A key challenge with understanding infectious disease spread in wildlife populations is that disease surveillance relies on imperfect sampling. Testing relies on the capture of live animals, or dead animals recovered from the field. Despite the use of systematic capture-mark-recapture (CMR) sampling—employed quarterly in WP—not all badgers are captured at each time point and capture rates vary seasonally. Furthermore, when most animals die their carcasses are not recovered, leading to considerable missing data and uncertainty regarding the number of badgers alive at any given time point. For endemic diseases, transmission dynamics depend on the mortality process, which for wild mammals must incorporate age-dependent mortality risks [13]. At WP, captured badgers are tested for bTB using a suite of different diagnostic tests: for the detection of antibodies (Brock, Dual Path Platform [DPP] and StatPak), a cellular immune response (the *γ*-interferon [*γ*-IFN] test), or the bacteria itself (microbiological culture). Diagnostic test performance (i.e. sensitivity and specificity) varies with the type of test, and test usage varied during the study period, with some being phased out (e.g.Brock/StatPak) and replaced by others (e.g. *γ*-IFN, DPP).

To tackle these challenges we developed a framework that simultaneously captures spatio-temporal disease dynamics, the CMR / mortality processes, and diagnostic test performance at the individual-level. Performing robust statistical inference on such a model is highly challenging, since model dynamics rely on a set of (often high-dimensional) unobserved epidemiological states that must be inferred as part of the model fitting process. Gold-standard approaches to this problem are computationally expensive [28, 29], and common approximation approaches are usually less suitable when considering highly-resolved (such as individual-level) data [30]. Instead we utilise a recent advance in Bayesian modelling known as the individual forward filtering backward sampling (iFFBS) algorithm [31], which provides an efficient and flexible inference methodology. We used this to fit a discrete-time

Susceptible-Exposed-Infectious-Dead (*SEID*) compartmental model of disease transmission to individual-level data from 2,391 badgers across 34 core social-groups captured during the WP study between 1980–2020; the model fitting in a few hours on a desktop machine.

## Materials and methods

The data are collected and curated by the Animal and Plant Health Agency on behalf of the Department for the Environment, Food and Rural Affairs, and are described in Section A1 of S1 Appendix. For data access requests, please contact. Badgers were trapped as part of a wider ongoing study at Woodchester Park under a Natural England Science and Conservation licence. Clinical samples were collected by experienced personnel holding relevant Personal Home Office licences and acting under a Home Office Project licence. All animals were examined and sampled under the supervision of a Named Veterinary Surgeon and a Named Animal Care and Welfare Officer. All work with live animals at Woodchester Park is subject to approval and periodic review by the Animal and Plant Health Agency Animal Welfare and Ethical Review Board.

We model transmission through a discrete-time compartmental model, defined over quarterly periods, where at any point in time individuals are either susceptible-to-infection (*S*), infected but not yet infectious (i.e. ‘exposed’—*E*), infected and infectious (*I*) or dead (*D*). Infected individuals enter the *E* class, and can then transition to the *I* class. However individuals can transition to the *D* class from any other class according to the mortality process (described below). The transmission rate in a given social group *s* at time *t* depends on the number of susceptibles, infectives and the total social group size, with a scaling parameter *q* used to estimate the degree of density vs. frequency dependence in transmission [32]. We assume that the social group structure is known, and that each badger had only moved to a new social group if it was captured there. Individuals born during the study period are assumed to be susceptible at birth, and those born before the monitoring period begins in any social group are given a non-zero probability of being in either the *S, E* or *I* states once monitoring commenced.

The mortality process in any mammal population is age-dependent, and conceptually a bathtub-shaped hazard function would be appropriate [33], where high early-life mortality risk drops to a constant level during mid-life and then increases exponentially in later-life. In wild badgers early-life mortality is often hidden, since deaths in newborns occur underground and so are never observed [13]. Instead, we model the *conditional* mortality distribution given survival to first capture as a Gompertz-Makeham distribution [34, 35]. We assume that badgers surviving this period of early-life mortality can be captured according to seasonally-varying (quarterly) capture probabilities. As such, mortality curves are typically right-censored at the last capture time for most badgers. The capture process is important for identifying the mortality parameters, since despite rarely recovering dead badgers the probability that an animal is alive but evades capture at successive time points quickly becomes negligible unless the capture probabilities are very small [13].

Environmental contamination (e.g. from bacteria shed in urine and faeces) may be a major source of bTB infection in badgers and cattle [36], and studies have shown that external sources of infection can contribute to bTB persistence in badger populations [37]. Several studies analyse the spatial organisation of badger populations and its implications for disease transmission at different spatial scales [38–43]. We consider social-group specific background rates-of-infection, to represent indirect transmission between badgers and infection from other sources such as cattle. We do not consider a direct between-social group badger-to-badger transmission process, outside of the explicit movements of individual animals between groups captured in the data.

Diagnostic tests are assumed conditionally independent *given the infection status* of each animal, and thus the observation process can be governed by the sensitivities and specificities of the different tests. This work extends ideas in [12] to incorporate a fully mechanistic model of disease transmission and progression, meaning that we are able to infer key quantities-of-interest that result from the mechanistic model, such as individual effective reproduction numbers. Since there is no recovered state in this system, the mortality process limits the length of the infectious periods for individual badgers. We also fit to a much longer time period (1980–2020); incorporating two additional types of test (Brock and DPP) and requiring a change-point for the Brock test sensitivities and specificities. Mathematical details of the model and parameters are given in Section A2 of S1 Appendix. We used vague or weakly informative prior distributions for the parameters (see Section A2.3 in S1 Appendix).

The iFFBS algorithm operates within a data-augmented Markov chain Monte Carlo (MCMC) routine. Standard FFBS approaches scale exponentially in the number of individuals, but the iFFBS algorithm (in certain situations) can be made to scale linearly with the number of individuals, substantially reducing computational overheads. We employed a combination of Hamiltonian Monte Carlo, Gibbs Sampling and Metropolis-Hastings updates for the parameters (see Section A4 in S1 Appendix).

Posterior predictive samples for the test results from individual badgers can be easily obtained and aggregated to any spatio-temporal level, enabling an assessment of model fit against the observed data. Similarly, posterior distributions for the relative contribution of direct badger-to-badger transmission on any new infection can be calculated through the ratio of the badger-to-badger transmission rate over the total transmission rate in the corresponding social group at each given infection time, which can again be aggregated to any desired spatio-temporal resolution (see Section A6 in S1 Appendix).

To quantify superspreading, we estimate individual effective reproduction numbers, *R*_*i*_, from which a population-level effective reproduction number, *R*, can be derived [16–18]. We define *R*_*i*_ to be the *expected number of secondary infections caused by an infected individual i across its lifetime*. We choose to calculate *R*_*i*_ only for animals that have died, and so each estimate adjusts for the latent and infectivity processes across the lifetime of each animal, and automatically accounts for the social-group specific background infection rates and changes in group sizes over time (and also any movements of badgers between social groups). Our *R*_*i*_ estimator is a variation of the likelihood-based approach of [16], in which the authors consider pairs of cases and use the serial (or generation) interval distribution (the time from symptom onset in a primary case to symptom onset in a secondary case) to derive a relative likelihood that a given case *j* was infected by case *i*. The individual-level reproduction number for an individual *i* is then a weighted sum of these relative likelihoods over all future cases.The generation-time interval is non-trivial to define here, since the infectious period is governed by the age-dependent mortality distribution. However, since we sample the hidden states for each individual, it is possible to derive a relative likelihood that a given individual was infected by any source (including the background infection rate), which can then be used in a similar way to produce estimates of *R*_*i*_. Instead of being limited to the onset of clinical signs, we integrate directly over the time-since-infection for each individual. For full details see Section A5 in S1 Appendix.

Code used to fit this specific model is available at https://github.com/evandrokonzen/WP bTB code and through the Environmental Information Data Centre repository (https://doi.org/10.5285/fe0f6bd7-ffd2-4e21-8c84-493cf4f3080d).

## Results

A total of 25,000 iterations ran in a matter of hours, and were sufficient to achieve good mixing of parameters (see Figs A1 and A2 in S1 Appendix) and event times. We ran two chains, discarded the first 5,000 as burn-in and then thinned the remaining samples to give 5,000 iterations in total across both chains. Comparison of posterior predictive outputs against the observed data shows that the fitted model successfully captures the dynamics of test-positive and test-negative animals across the whole population for all test types over the complete 40+ year time period (Fig 1). Analogous plots for each social group (Section A7.3 in S1 Appendix) show similarly good fits, justifying the use of the model to examine key drivers of disease transmission in WP.

**Fig 1.**
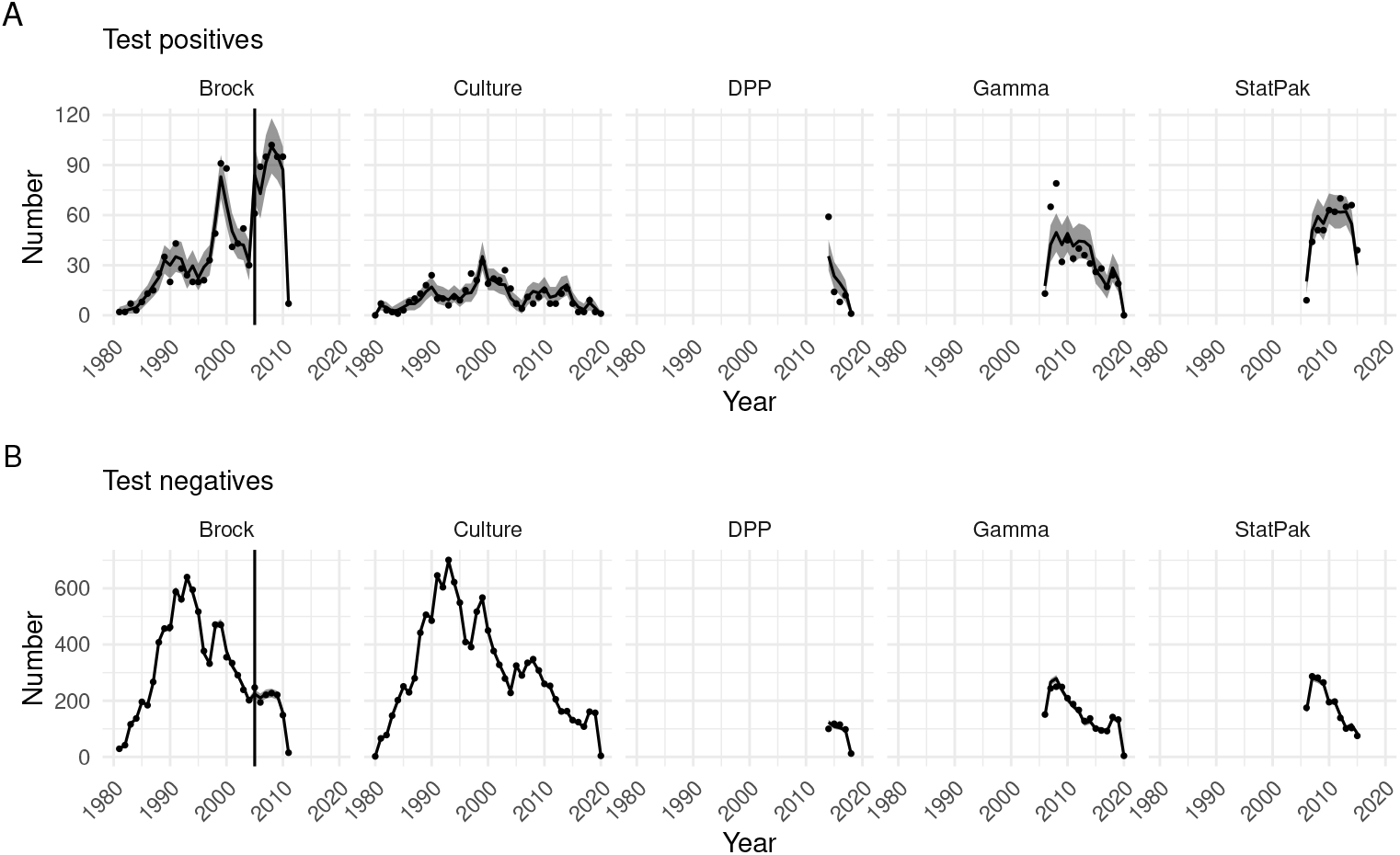
Posterior predictive summaries for the numbers of test positive and negative individuals over time. Plots show number of (A) test-positive and (B) test-negative animals for each test in each year against the observed data. Posterior means are indicated by black lines and 95% credible intervals by grey ribbons. Data are represented by black points. The vertical black line in the Brock panel corresponds to the posterior predictive mean of the changepoint for this test.

### Reproduction numbers

Posterior summaries for the individual- and population-level reproduction numbers (Fig 2) reveal substantial heterogeneity in reproduction numbers between individuals: some badgers have *R*_*i*_ *≈* 8, whereas most have *R*_*i*_ *<* 1. Despite some animals having high individual reproduction numbers, the population effective *R* is on average *<* 1 (Fig 2B) with a posterior mean estimate of 0.66 (95% credible interval [CI]: 0.59–0.72). To explore key factors that might contribute to the estimated heterogeneity in *R*_*i*_, we plot the posterior mean *R*_*i*_ values against a set of other individual-level summary measures: the posterior mean infectious period, the average number of social groups a badger belonged to, and the average size of social groups a badger belonged to (Fig 3). Heterogeneity in the duration of the infectious period due to variations in infection time and survival seems to be the main (although not sole) driver of these differences (Fig 3A). Since *R*_*i*_ values do not associate strongly with the average number of social groups that a badger has been associated with, or with average group sizes (Figs 3B–D), it is most likely that on average badgers infected at a younger age simply have more opportunity for onwards transmission. We also compare the posterior mean probability of infection against the posterior mean *R*_*i*_ values across individuals and show that superspreading is not just a function of the probability of infection (S1 Fig).

**Fig 2.**
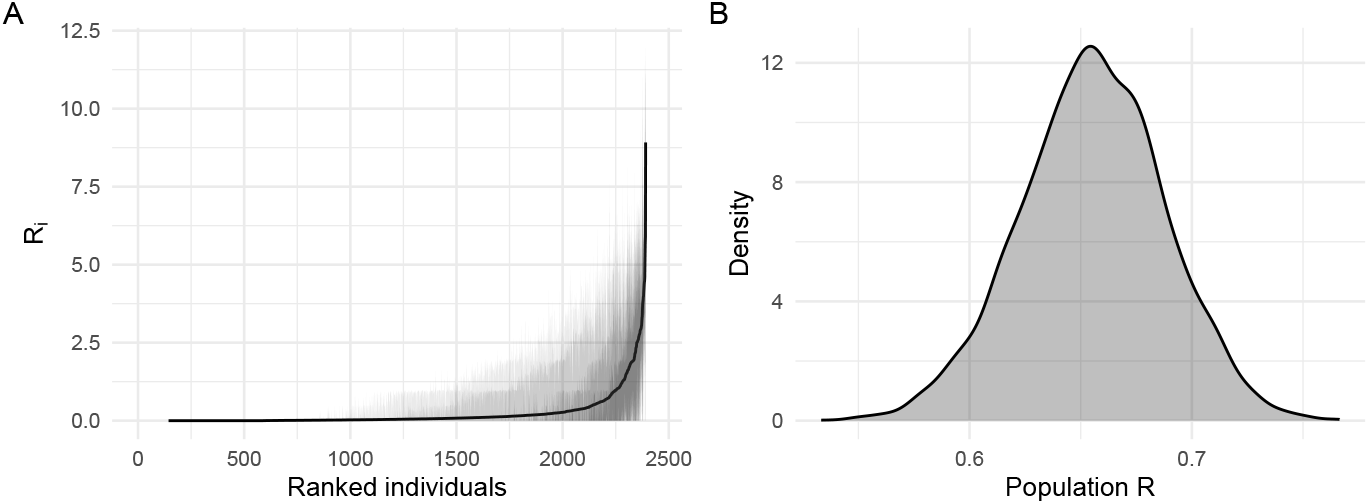
Posterior summaries of reproduction numbers. (A) Posterior means and credible intervals (50% and 95%) for the individual reproduction number for all individuals, ranked in increasing order of posterior means, and (B) posterior distribution for the population effective reproduction number.

**Fig 3.**
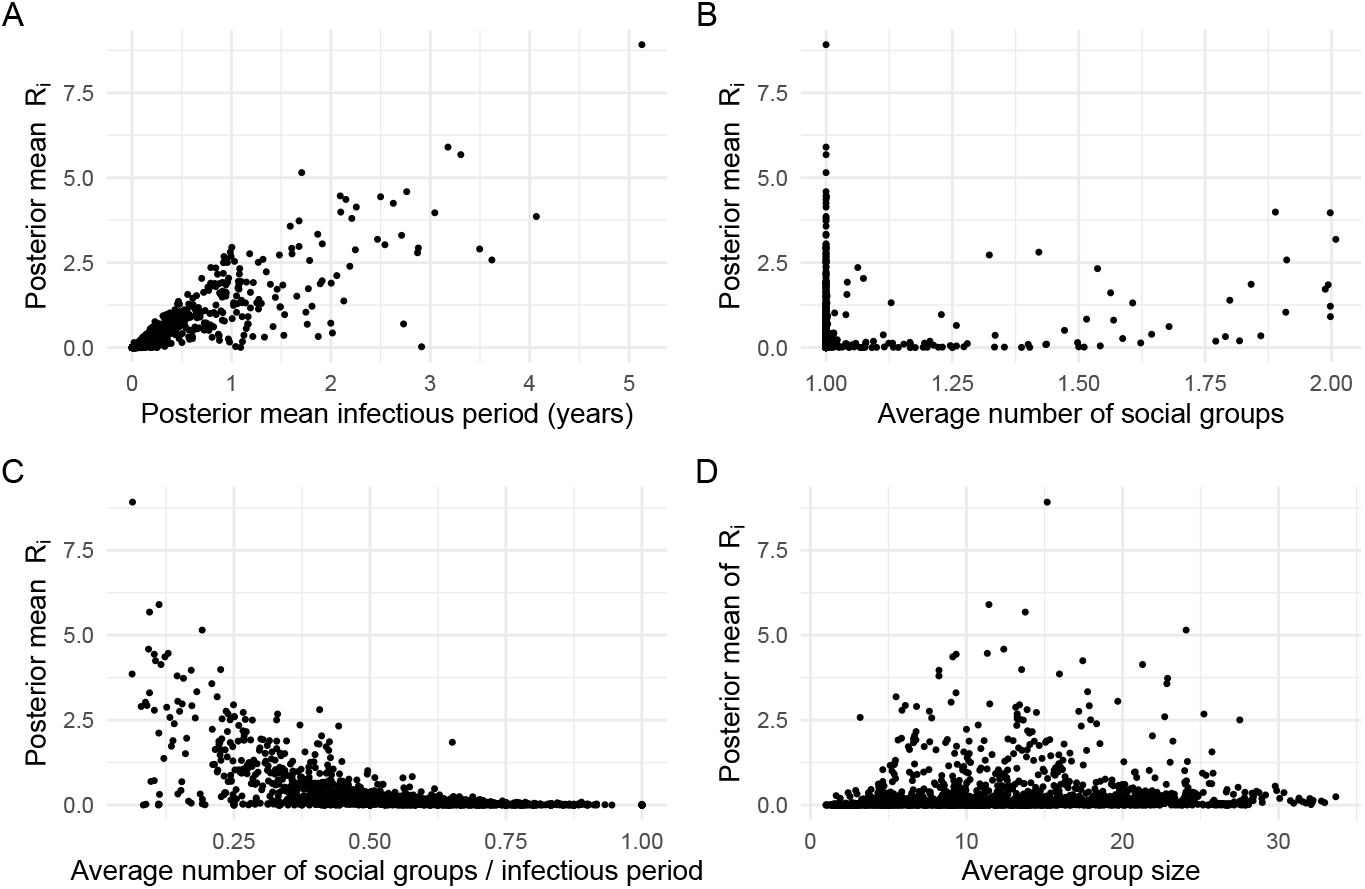
Posterior mean estimates of individual reproduction numbers against other factors. Plots show the posterior mean *R*_*i*_ against (A) posterior mean duration of the infectious period; (B) average number of social groups an individual belonged to; (C) average number of social groups an individual belonged to divided by the duration of the infectious period; (D) the average size of social groups that the individual belonged to.

### Hidden burden of infection

We also estimate the probability of infection/infectivity of any badger at any given time, and the corresponding event times (e.g. S2 Fig), accounting not only for the diagnostic test history for each animal, but also information from other badgers as captured through the transmission model. These estimates can be aggregated up to higher spatial levels, such as the social-group or population levels. For example, Fig 4A shows the predicted number of individuals in each epidemiological state across the whole population over the study period, giving an estimate of the overall hidden burden of infection [44]. Analogous plots for each social group are shown in S3 Fig and S4 Fig. The model predicts relatively low levels of underlying infection in the population over the study period, however, there is a steep decline in population size from about 1993 onwards, with the proportion of infected badgers increasing towards the end of the study period. We predict that on average 72% (95% CI: 65%–78%) of new infections can be attributed to direct badger-to-badger transmission, with the rest attributed to the background rate-of-infection terms (Fig 4B). In addition to transmission arising from other species (e.g. cattle), the background infection risk can also encompass indirect transmission from badgers, such as from environmental contamination, and hence the 72% estimate represents a lower bound for the relative rates of *all* badger-to-badger transmission in WP.

**Fig 4.**
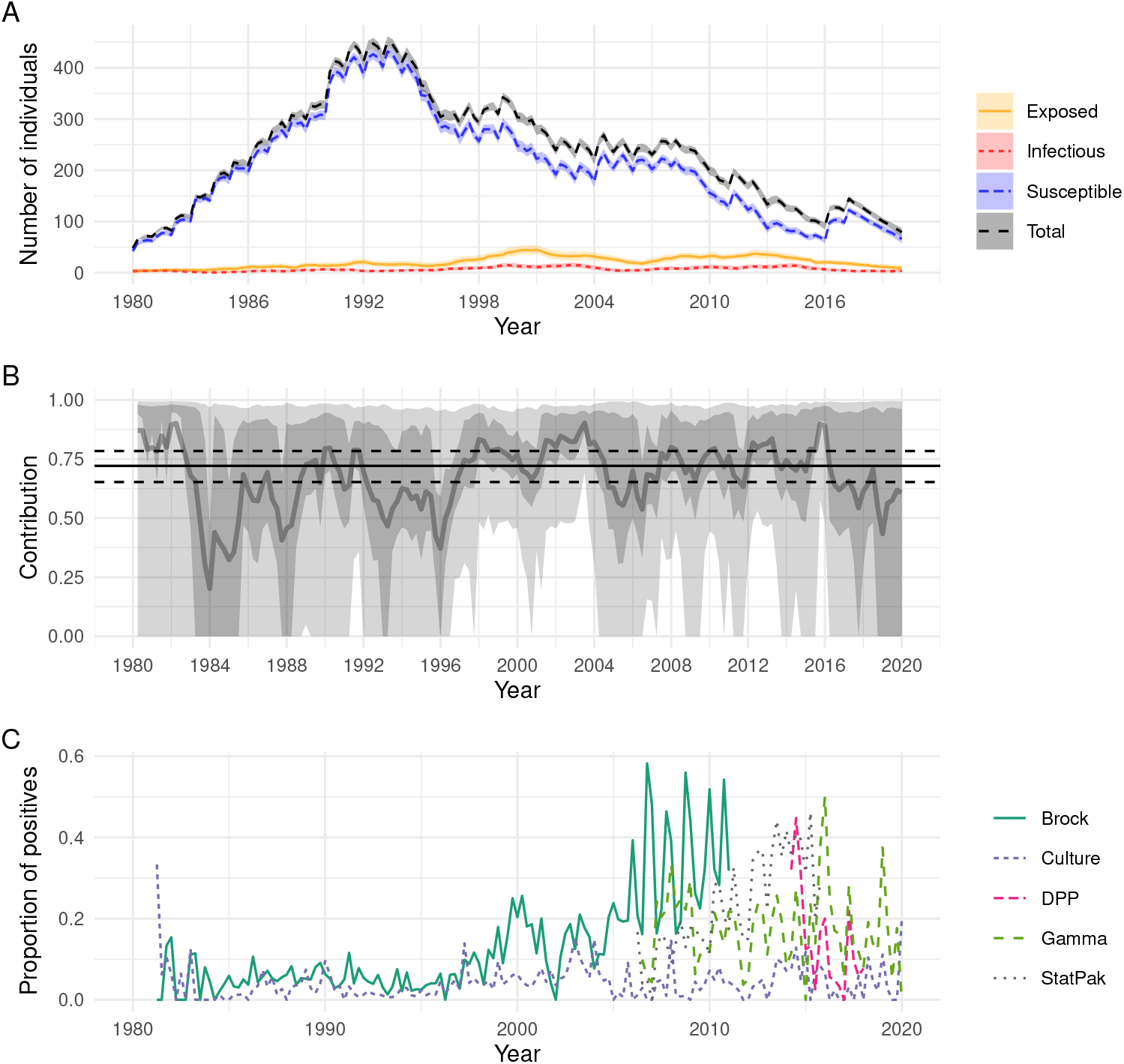
Posterior predictive summaries for aspects of the underlying epidemic. (A) Posterior predictive means and 95% prediction intervals for the number of individuals in each epidemiological state over time; (B) posterior predictive means and credible intervals (50% and 95%) for the relative contribution of badger-to-badger transmission vs. background rates-of-infection on new infections over time (the solid black line is the average contribution over all time points, with the dashed lines giving the corresponding 95% credible interval); (C) the observed proportion of positive test results over time for each test.

### Epidemiological parameters

Posterior summaries for all epidemiological, CMR, mortality and diagnostic test parameters are shown in Table 1, and Figs A3–A4 and Table A1 in S1 Appendix. We obtain an estimate of the average (discrete-time) latent period (the time between infection and infectiousness) of 3.7 (95% CI: 2.9–4.7) years. We also obtain estimates for the infectious period distribution, which is derived from the age-dependent mortality curves and the inferred infection times. To understand the transmission potential of individual badgers, we define an *effective infectious period* distribution as the time between when an animal becomes *exposed* (i.e. when they enter the *E* class) and when they die. We then define the *infectious period* as the time between when an animal becomes *infectious* (i.e. when they enter the *I* class) and when they die, here equivalent to the conditional survival distribution given survival beyond infectiousness. Since we simulate the time-of-infectiousness and the time-of-death for each animal, we can generate empirical estimates for these distributions that average over the unobserved event times (Fig 5). It is clear that a large proportion of badgers die before becoming infectious, and so have an *effective* infectious period of zero (Fig 5A); shown even more clearly if averaged over the population (Fig 5B). Considering the infectious period distribution *conditional on being infectious*, then we obtain a posterior mean of 1.4 (95% CI: 0.25–4) years averaged across the population (Fig 5C).

**Table 1.** Posterior means and 95% credible intervals for key model parameters. Results given to 2 significant figures (4 s.f. for changepoint). For full details of the model structures and parameters, see Section A2 in S1 Appendix.

| Type | Parameter | Mean | 95% CI |
| --- | --- | --- | --- |
| Transmission | $\beta$ | 0.019 | (0.009, 0.037) |
| | $q$ | 0.59 | (0.23, 0.91) |
| | $\tau$ (years) | 3.8 | (3, 4.8) |
| Mortality | $a$ | 0.036 | (0.032, 0.039) |
| | $b$ | 0.033 | (0.029, 0.037) |
| | $c$ | 0.0016 | (0.000034, 0.0056) |
| Sensitivity ($E$) | Brock1 | 0.56 | (0.49, 0.63) |
|  | Brock2 | 0.73 | (0.65, 0.8) |
|  | Culture | 0.073 | (0.05, 0.098) |
|  | DPP | 0.41 | (0.31, 0.53) |
|  | Gamma | 0.47 | (0.41, 0.52) |
|  | StatPak | 0.82 | (0.75, 0.88) |
| Sensitivity ($I$) | Brock1 | 0.91 | (0.84, 0.96) |
|  | Brock2 | 0.93 | (0.83, 0.99) |
|  | Culture | 0.8 | (0.73, 0.86) |
|  | DPP | 0.92 | (0.76, 1) |
|  | Gamma | 0.85 | (0.76, 0.93) |
|  | StatPak | 0.95 | (0.89, 0.99) |
| Specificity | Brock1 | 0.98 | (0.98, 0.99) |
|  | Brock2 | 0.78 | (0.76, 0.8) |
|  | Culture | 0.99 | (0.99, 1) |
|  | DPP | 0.94 | (0.92, 0.97) |
|  | Gamma | 0.92 | (0.91, 0.93) |
|  | StatPak | 0.94 | (0.93, 0.96) |
| Capture Probability | Winter | 0.18 | (0.17, 0.18) |
|  | Spring | 0.36 | (0.35, 0.37) |
|  | Summer | 0.46 | (0.45, 0.47) |
|  | Autumn | 0.28 | (0.27, 0.28) |
| Changepoint | $\xi$ | 2005 | (2005, 2005) |

**Fig 5.**
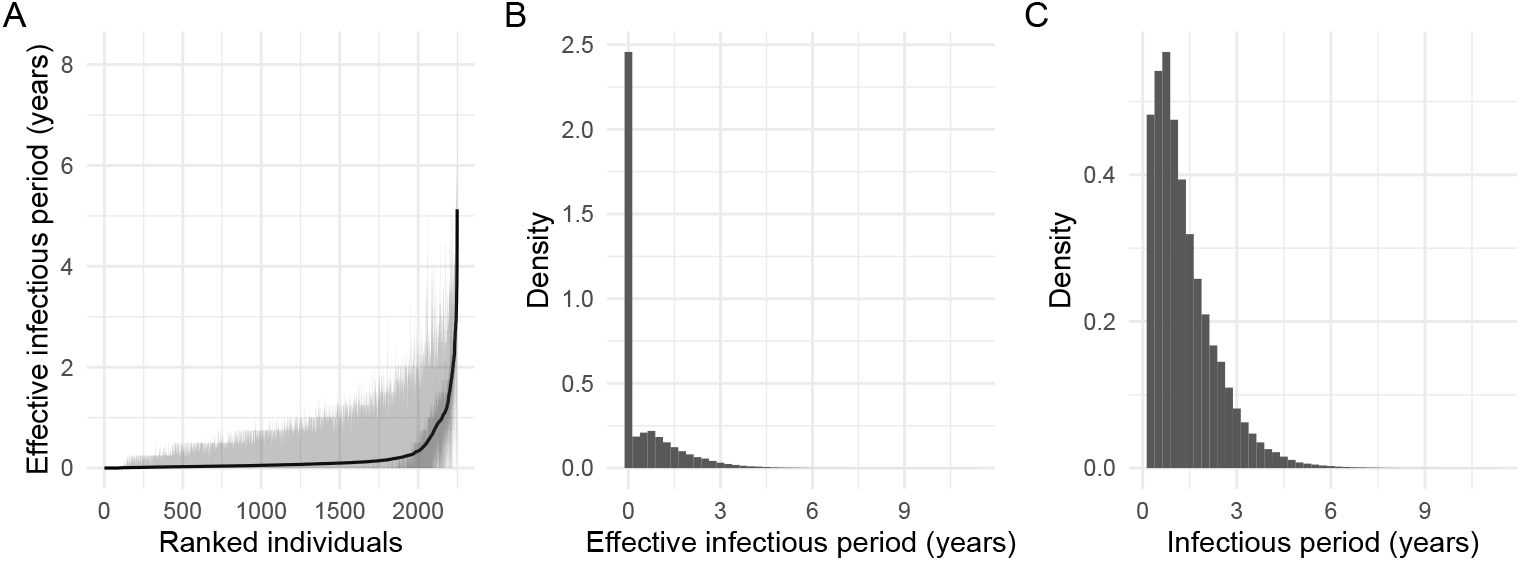
Posterior summaries for the length of the infectious periods. (A) Posterior means and credible intervals (50% and 95%) for the individual effective infectious period distribution across all individuals, ranked in increasing order of posterior means; (B) posterior distribution for the population-averaged *effective* infectious period distribution; (C) posterior distribution for the population-averaged infectious period distribution.

Transmission can be considered as lying on a continuum between density- and frequency-dependent, captured here by a parameter *q*, which mediates the extent to which contact rates are governed by badger density [32]. We infer the posterior mean for *q* to be 0.59 (95% CI: 0.23–0.91), which provides evidence consistent with the transmission process being somewhere in-between frequency- and density-dependent (defined when *q* = 1 and *q* = 0 respectively; [32, 44]). Following [44], our choice of a *U* (0, 1) prior distribution stops *q* from being exactly 0 or 1, but the area of high posterior density is nevertheless away from either of these two extremes.

### Mortality and capture processes

We estimate a median survival time of 3.6 (95% CI: 3.5–3.8) years, with posterior predictive plots of the survival and hazard functions shown in S5 Fig. We estimate seasonal differences in capture probabilities, ranging from 0.18 (95% CI:0.17–0.18) in the winter, to 0.46 (95% CI: 0.45–0.47) in the summer (Table 1).

### Diagnostic test performance

We estimate diagnostic test sensitivities and specificities relative to infection, rather than to a proxy measure such as visible lesions [12, 44]. In the first version of our model we assumed that each of the five tests had constant sensitivity and specificity values over time. We obtained posterior mean estimates for the sensitivities (for infectious animals) of: Brock 0.88 (95% CI: 0.84–0.91), Culture 0.32 (0.29–0.35), StatPak 0.93 (0.89–0.96), *γ*-IFN 0.57 (0.53–0.62) and DPP 0.63 (0.53–0.73). The sensitivity of the *γ*-IFN test was much lower than expected from cattle studies [45], although previous estimates in badgers are more variable: [12] estimated a *γ*-IFN sensitivity of about 0.44, whereas [10] estimated a value of 0.799; although the former did not differentiate between latently infected and infectious badgers (which may drive the estimate to be lower if sensitivity is lower during the latent period), and the latter used a different method but had a more informative prior distribution with a median sensitivity of 0.796.

This original low sensitivity estimate for the *γ*-IFN test led us to explore other possible explanations. Plotting the proportion of test positive results over time for the five tests, we see that from about 2005 onwards the proportion of test-positives for the Brock test increases rapidly, whilst the other tests (notably culture) drop or remain constant (Fig 4C). The exception is the StatPak (an antibody test like the Brock), and similar inconsistencies with the other tests have previously been noted [46].

Consequently, we developed our current model, where the Brock test sensitivity and specificity were permitted to change at some changepoint. Following this modification the posterior distribution for the changepoint is estimated to be between the final quarter of 2004 and the first two quarters of 2005 (Table 1), consistent with the data in Fig 4C. Consequently the pre- and post-changepoint Brock test sensitivities (denoted Brock1 and Brock2 respectively) are both high (0.91 and 0.93), but the specificity of the Brock post-changepoint drops to 0.78 (from 0.98 pre-changepoint). The inclusion of the changepoint also results in the *γ*-IFN sensitivity estimate rising to about 0.85, which is more consistent with earlier literature (except [12] as discussed above). Model fits, parameter estimates and predictive plots for the initial (non-changepoint) model are shown in the Section A7.4 in S1 Appendix. We note that the model fit was not as good for this model (Fig A5 in S1 Appendix).

## Discussion

We have inferred a suite of epidemiological parameters describing the epidemiology of bTB in an undisturbed population of badgers, monitored for several decades. Our key findings include characterising heterogeneity in the latent and infectious periods of individual hosts, and most importantly heterogeneity among individuals in their individual effective reproduction numbers. Integrated across the whole population, we infer the population-level reproduction number to be credibly below one. We discuss these key findings, and the importance of other parameter inferences, in turn.

Heterogeneity in individual reproduction numbers implies the presence of superspreader badgers, a result that emerges from the stochastic model after fitting it to the observed data [19], and highlights the importance of accounting for stochastic dynamics when modelling disease outbreaks in small- to medium-scale populations. The existence of superspreaders raises interesting questions around the mechanisms contributing to the endemicity of bTB in WP. Periodic reintroductions from external sources and/or persistent environmental contamination, such as shedding, could serve to maintain infection in the population—captured here through the background rate-of-infection terms. Our model is based on a subset of badger social groups, and reintroduction of infection into this meta-population could also occur through interactions with unmodelled neighbouring social groups in the wider landscape. With currently available data we are not able to disentangle these mechanisms, but since the population-level *R* is estimated to be *<* 1 with a high posterior probability, it seems likely that a suitable management strategy such as vaccination or selective removal of superspreaders would be an efficient means of driving infection rates down, since transmission events are more likely to occur by direct badger-to-badger transmission, predominantly from superspreading individuals. Social-group specific predictions (e.g. S3 Fig and S4 Fig) could be useful tools to guide social-group or even sett-specific surveillance efforts. A key question for further research is whether reintroduction of infection from external sources could be sufficient to maintain infection in the face of this type of management strategy within this study area.

Other modelling studies have assumed or estimated slightly higher *basic* reproduction numbers (*R*_0_) for bTB in cattle and wildlife populations of just above 1 [7, 47–49], though estimates are often made using simpler population-level (often deterministic) models with constant mortality assumptions. A recent paper estimated the within- and between-species reproduction numbers for cattle and badgers from data in Kilkenny, Ireland, using an explicit spatial connectivity structure between badger social groups and cattle farms with an environmental decay term to capture persistence of the pathogen in the environment [9] (see also [50]). Their analysis suggested that environmental transmission can play a major role in disease transmission, and determined what relative badger-to-cattle densities and badger vaccination coverage can cause the basic reproduction number to be above or below one in different cattle herds. Nevertheless, these previous estimates are not directly comparable to our population-level effective *R*, since *R*_0_ characterises the expected number of secondary infections per primary infection introduced to a *fully susceptible population*. In contrast, our population-level *R* characterises the posterior expected number of secondary infections per primary infection that has occurred within our study population, which will be lower than *R*_0_ due to factors such as varying numbers of susceptible badgers over time, the meta-population structure or changes in social group sizes. Our estimate also adjusts for the background risk-of-infection (which if ignored will assign higher weight to badger-to-badger interactions) and age-dependent mortality. This study adds to the literature suggesting that the population-level reproduction number for badgers is likely to be low [1, 7].

We have estimated the burden of infection to be relatively low throughout the study period, but with an increase in prevalence coinciding with a decline in host population size. We estimate direct badger-to-badger transmission to account for 72% of new infections, but we consider this a lower bound for all badger-to-badger transmission, and future work will attempt to disentangle some of these competing processes.

Historic models of bTB in badgers produced estimates for the latent period of around 1–1.5 years [51], but these were estimated from population-level deterministic models with constant mortality rates. Other approaches estimated shorter latent periods of between 95–158 days, but these were estimated in a laboratory setting [52]. In contrast we estimate a longer average latent period of 3.7 years. In cattle, estimates vary more widely; typical assumptions are between 6–20 months [44], but some estimates are much longer (*>* 11 years) [50]. Our average latent period estimate is similar to our estimated median survival time for badgers of 3.6 years (see also [13]), which is slightly longer than earlier estimates for expected life-expectancy of between 2.2–2.9 years assuming constant mortality rates [53]. Our survival estimates are conditional on *at least one capture event* (since early-life mortality events occurring in underground burrows (setts) are unobserved), and hence are expected to be slightly longer than the true survival time. Correcting for this bias is the focus of future work.

We also produce posterior distributions for the length of the infectious period for each badger, which accounts for the infection process and the age-dependent mortality. When averaged across the population, these suggest a high probability of infected badgers dying before they become infectious; introducing natural heterogeneity in the degree to which individual badgers can transmit bTB. In comparison, [9] assume a mean infectious period of 1 year, with a range between 0.32–3.6 years (from [52, 53]). In cattle, accurate estimation of the latent and infectious periods is challenging due to censoring and selection induced by the routine slaughter of test-positives, but these challenges should be mitigated here since no artificial control measures were implemented in this population over the study period.

We infer the transmission process for bTB to lie in-between the extremes of density- and frequency-dependence. Frequency-dependence is a common assumption for badger bTB models [9, 54, 55], and our estimated *q* is not dissimilar from some earlier cattle studies [44]. This result suggests that contact rates will increase with badger density at low densities, and then begin to saturate at higher densities [32]. This also has implications for disease control strategies, because under frequency-dependent transmission, population-reduction is a less viable means of disease eradication [56].

Our finding of a change-point in the sensitivity and specificity of the Brock diagnostic test also suggests some change in the Brock test itself or how it was delivered after 2005, which is an ongoing area of investigation. The clear reduction in Brock test specificity in later years also explains another phenomenon in the observed data, which is that some individual badgers consistently tested positive by the Brock test but negative by the other tests (see also [46]).

Estimates of the hidden epidemiological states change dramatically when the change in Brock sensitivity and specificity is included, with the original model showing much higher levels of underlying infection than the changepoint model, with higher rates of badger-to-badger transmission. We note that both models suggest the presence of superspreader badgers with a population-level *R <* 1. However, the changepoint model is preferred here, for several reasons: the model fits were better; more credible estimates for the other test sensitivities and specificities were obtained; and the model acknowledges previously highlighted concerns with the Brock test specificity in later years.

### Conclusions and future work

To summarise, we have developed an individual-level stochastic meta-population compartmental model of bTB in badgers and fitted this to a dataset derived from an intensive long-term capture-mark-recapture study using an iFFBS algorithm. The model fitted in a few hours, enabling us to conduct efficient Bayesian inference simultaneously on all epidemiological parameters, as well as those relating to age-dependent mortality, the CMR surveillance process and diagnostic test performance.

The model provides novel estimates of individual effective reproduction numbers, suggesting that a small proportion of infected badgers, or superspreaders, contribute disproportionately to the transmission potential of the disease. This is despite the population-level effective reproduction number being low, suggesting that management strategies could be improved by targeting these animals or associated social groups with interventions or more intensive surveillance. Given systematic surveillance data, the model produces predictions for the hidden epidemiological states of individual badgers over time, which can be aggregated to different spatio-temporal resolutions. This could be used to help target surveillance or management strategies since we can estimate the likelihood of individuals being infected or infectious, and the expected infectious period, accounting for the age-dependent mortality process and the infection times. These estimates are not just based on the diagnostic test history for individual animals, but also borrow weight from other animals through the transmission model structure.

Detailed predictive information like this is only feasible in a population with systematic surveillance, such as in Woodchester Park, but the iFFBS algorithm has demonstrated its value in modelling such data efficiently, and could be employed in other settings; for example the modelling of within-herd spread of bTB in livestock. Nevertheless, there are various areas that could be developed further. For example, exploring more nuanced infectivity-profile dependent sensitivity and specificity processes, rather than the coarse step-functions that we currently employ could be insightful about disease progression [57]. It may also be valuable to include individual-level covariate dependencies in different components of the model, for example, to explore sex-specific differences in mortality [13]. Also of interest is investigating the effect of infection on mortality risk [13, 47], or sex-/age-specific differences in transmission/susceptibility. With suitable data it would be of interest to integrate different choices of spatial connectivity structure and/or environmental transmission, or to deal with multiple species [9, 50]. We also propose further exploration of the characteristics of superspreader badgers, which may inform operational targeting. The insights described above have been gleaned from a single plausible model, but a key area of future work is to extend this framework to allow for systematic model comparison and/or model averaging, which could provide even more robust predictions and help disentangle complex model interactions [29, 58].

## Supporting information

**S1 Appendix**. Supporting Information.

**S1 Fig.**
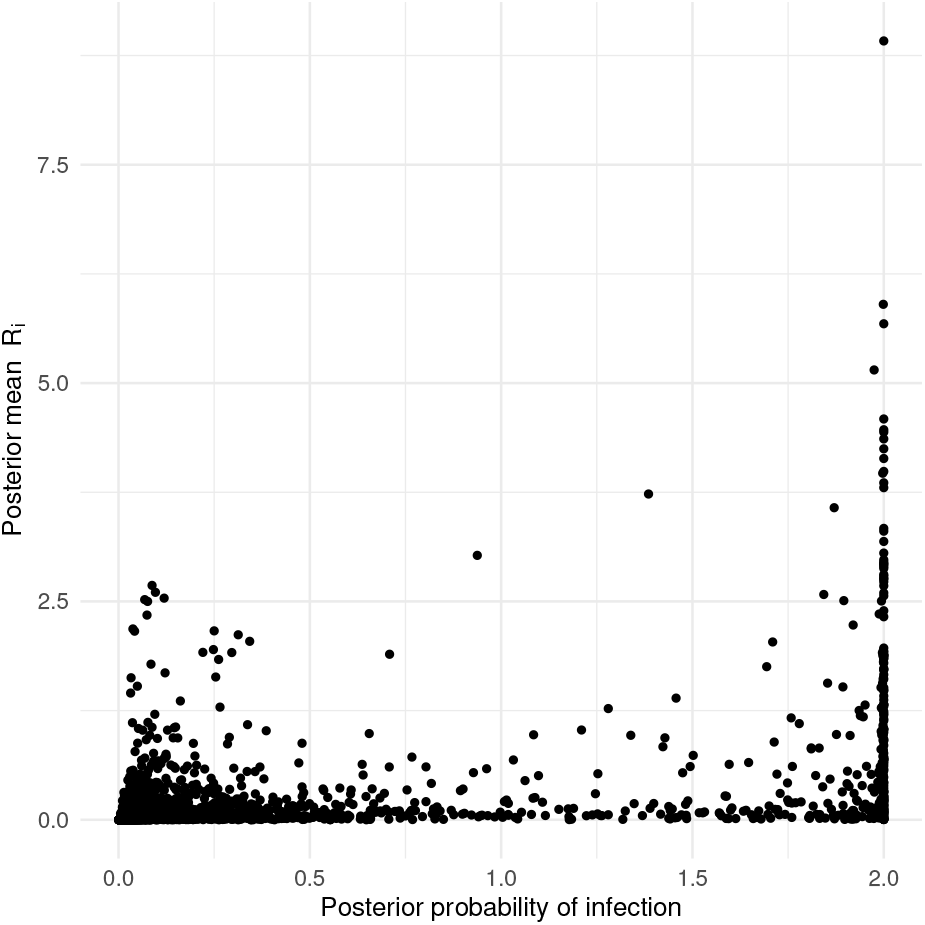
Posterior mean individual effective reproduction numbers against posterior probability of infection.

**S2 Fig.**
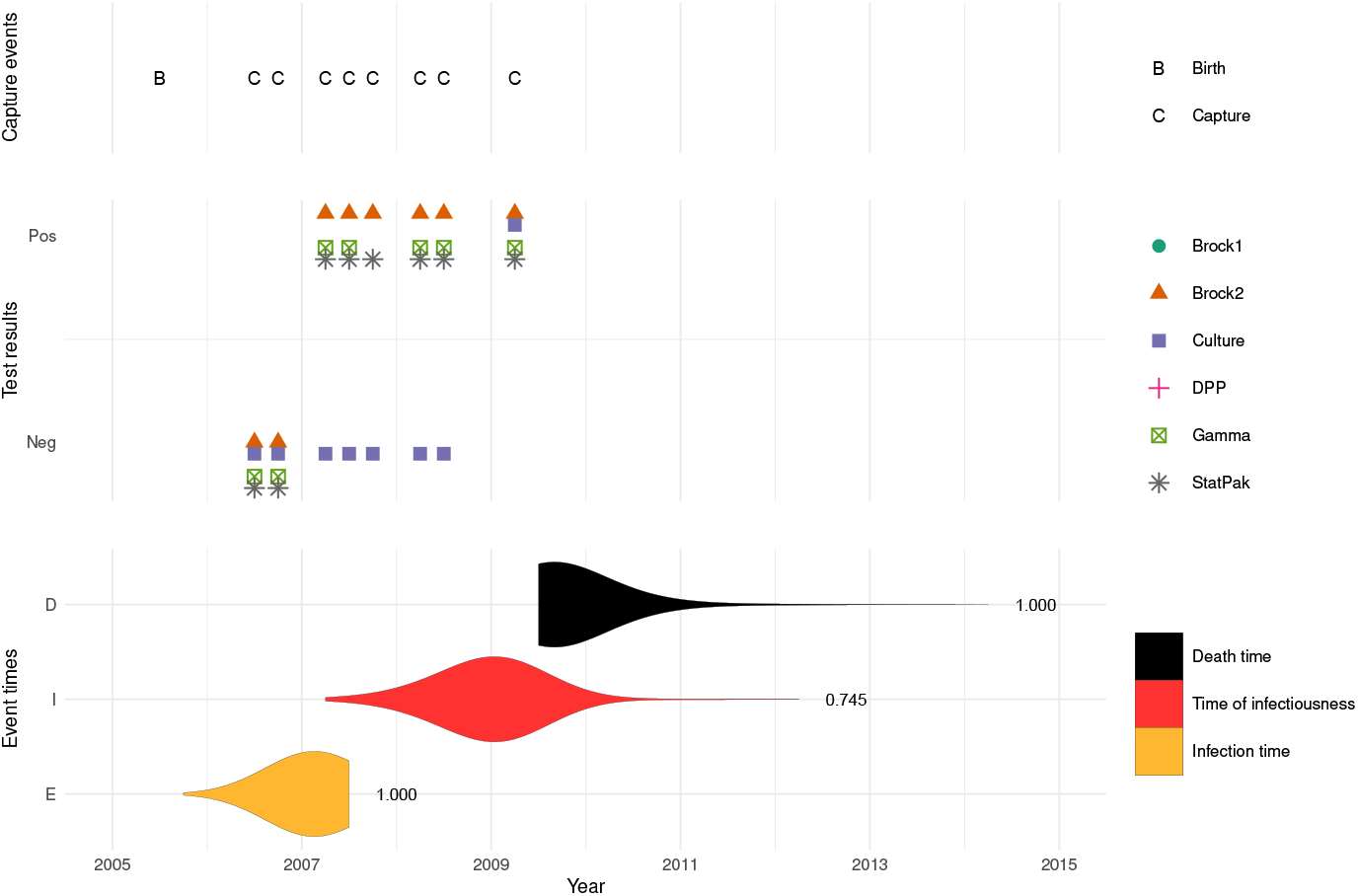
Capture history, test results and posterior distribution of event times for a single individual badger. Densities are the conditional posteriors for the event time given that the event occurred.

**S3 Fig.**
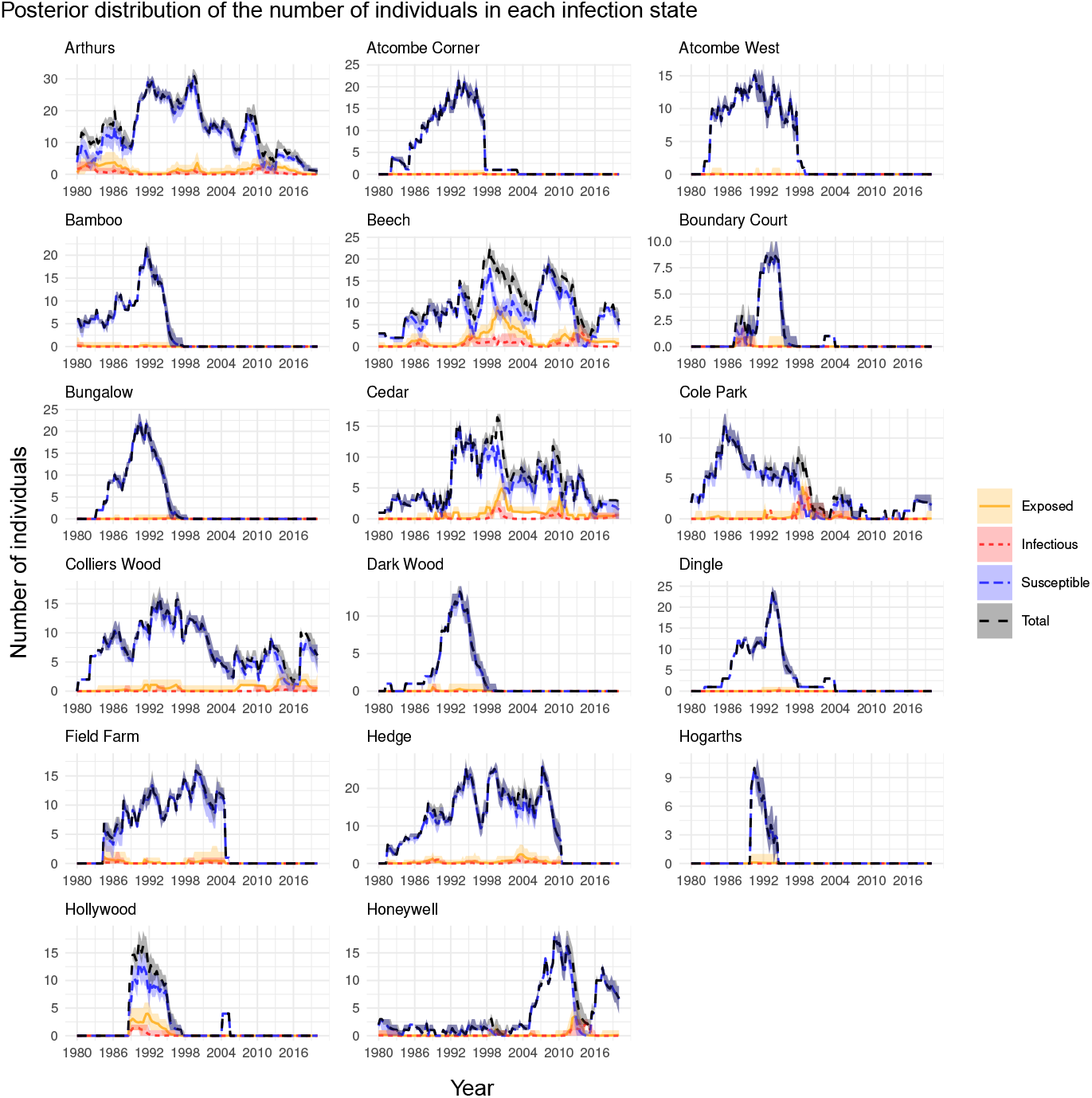
Posterior distribution (mean and 95% credible intervals) of the number of individual badgers in each infection state for social groups 1–17.

**S4 Fig.**
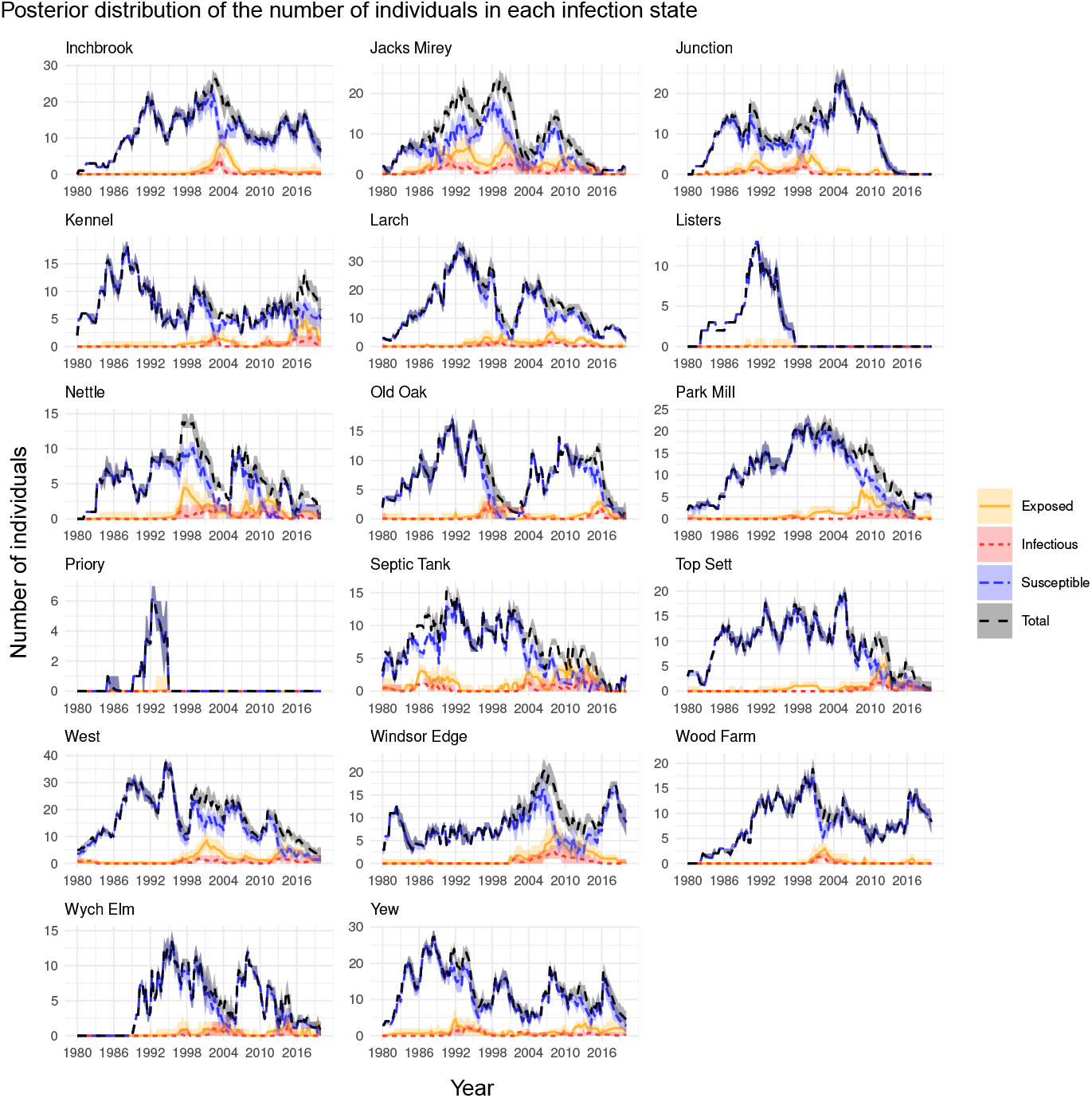
Posterior distribution (mean and 95% credible intervals) of the number of individual badgers in each infection state for social groups 18–34.

**S5 Fig.**
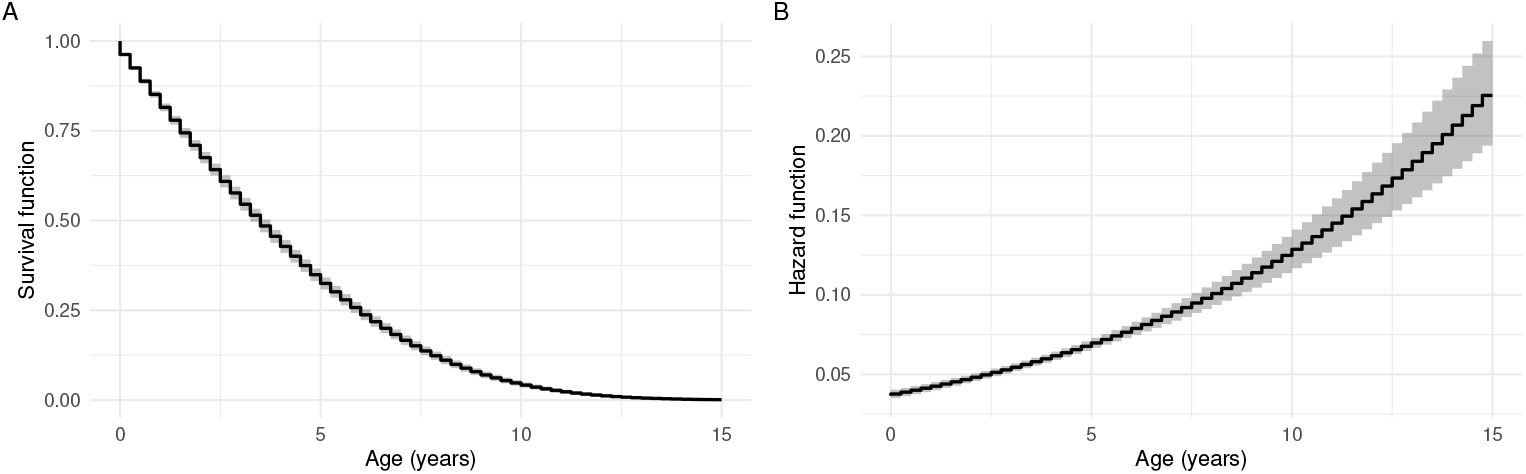
Posterior predictive plots for the mortality curves. (A) the survival function, and (B) the hazard function. Posterior means are indicated by black lines and 95% credible intervals by grey ribbons.

## Acknowledgments

We would like to thank Andrew Conlan for useful discussions. This work was funded by the Natural Environment Research Council grant number NE-V000616-1. We thank the many field and laboratory staff who have contributed to the collection of the unique Woodchester Park database. For the purpose of open access, the authors have applied a Creative Commons Attribution (CC BY) licence to any Author Accepted Manuscript version arising from this submission.

