## Supplementary Materials for "Efficient modelling of infectious diseases in wildlife: a case study of bovine tuberculosis in wild badgers"

### A1 Dataset and sampling period for hidden states

We use data from 34 social groups. The original dataset contains $m = 2751$ individuals, but we only use the individuals which, at their first capture, were found in a core social group. We discard 360 individuals (13.1%), out of which: 165 (6.0%) were never captured in any of the core social groups; 63 (2.3%) were initially captured in a non-core social group and moved to a core social group afterwards; 132 (4.8%) were first captured in a social group which was not included in the model.

At the known birth times (recorded at quarterly time intervals), we assume the badgers are not infected. We also assume that individuals belong to a social group until they are captured in a new social group. Badgers were sometimes captured and tested multiple times at the same quarter. All these test results are used and assumed to be independent given the hidden states.

Since trapping effort was consistent in some social groups at different times, each social group has its own monitoring period. In addition, we assume the monitoring period for an animal ends if it leaves the core area. We conduct inference on the hidden states of the animals only in their corresponding monitoring periods. However, trapping events occurring outside of the monitoring period are still used in the mortality component of the model.

For animals born during the monitoring period, we sample the infection states from their date of birth. For animals born beforehand, we assume they are susceptible, exposed, or infectious with probability $\nu_S$, $\nu_E$, and $\nu_I$, respectively, at the beginning of the monitoring period. Infection states of individual $i$ are sampled up to the end of the monitoring period ($T_i$).

### A2 Model

#### Notation

- $\Omega = \{S, E, I, D\}$: set of possible states (susceptible, exposed, infectious, dead)
- $N$: number of states (number of elements in $\Omega$)
- $m$: number of individuals
- $T$: number of time points of the discrete-time hidden state process
- $X_t^{[i]} \in \Omega$: hidden state variable for individual $i \in \{1, \ldots, m\}$ at time $t \in \{1, \ldots, T\}$
- $\boldsymbol{X}_t^{[1:m]} = \left(X_t^{[1]}, \ldots, X_t^{[m]}\right)'$: vector of infectious states of $m$ individuals at time $t$
- $\boldsymbol{X}_{1:t}^{[1:m]} = \left(\boldsymbol{X}_1^{[1:m]}, \ldots, \boldsymbol{X}_t^{[1:m]}\right)'$: set of $m$ individuals at times $1, \ldots, t$
- $\boldsymbol{X}_t^{[-i]}$: vector of all individuals, except $i$, at time $t$
- $W_t^{[i]}$: random binary variable for the capture process of individual $i$ at time $t$
- $\boldsymbol{Y}_t^{[i]} = (Y_{1t}^{[i]}, \ldots, Y_{Jt}^{[i]})$: random vector for the outcomes of the $J$ tests for individual $i$ at time $t$
- Data:
  - $w_t^{[i]}$: observed binary variable indicating whether individual $i$ is captured at time $t$: $w_t^{[i]} = 1$ if captured, and 0 otherwise
  - $o_{jt}^{[i]}$: observed binary variable indicating whether test $j$, $j = 1, \ldots, J$, was taken for individual $i$ at $t$: $o_{jt}^{[i]} = 1$ if the test was taken, and 0 otherwise
  - $\boldsymbol{o}_t^{[i]} = \left(o_{1t}^{[i]}, \ldots, o_{Jt}^{[i]}\right)'$: binary vector indicating whether individual $i$ was tested by each of the $J$ tests at time $t$
  - $y_{jt}^{[i]}$: observed result of test $j$, $j = 1, \ldots, J$, at time $t$ for individual $i$: $y_{jt}^{[i]} = 1$ if tested positive, and

0 if tested negative

- $\mathcal{I}_{i,t_i-1}$: set of individuals who could infect individual $i$ in the time period $[t_i - 1, t_i)$
- $\mathcal{M}_{i,t_i-1}$: set of individuals in the social group to which individual $i$ belongs at time $t_i - 1$ (including individual $i$ itself)
- $|\mathcal{X}|$: cardinality of the set $\mathcal{X}$
- $t_i^B, t_i^E, t_i^I$: times at which individual $i$ is born, becomes exposed, and becomes infectious, respectively
- $t_{0,i}$: starting time for the inference of hidden states of individual $i$. It is either the birth time or the start of the monitoring period, whichever is later
- $T_i$: end of monitoring period for individual $i$
- $T_i^{\text{LC}}$: last capture time of individual $i$
- $\psi_t^{[i]}$: probability that individual $i$ dies $[t-1, t)$ given it is alive at $t-1$.
- $\mathbb{1}_{\{A\}}$ indicator function that equals 1 if $A$ is true, and 0 otherwise
- $I_t^{[i]}(j, l)$: indicator function that equals 1 if individual $i$ is in state $j$ at time $t-1$ and in state $l$ at time $t$, and 0 otherwise.
- $\alpha_g$: background rate of infection in the $g$-th social group, $g = 1, \ldots, G$
- $\boldsymbol{\alpha} = (\alpha_1, \ldots, \alpha_G)'$: vector with all the background rates of infection
- $\alpha_{g_i(t_i-1)}$: background rate of infection of the social group that individual $i$ belongs to at time $t_i - 1$
- $\lambda$: parameter used in the hierarchical shrinkage prior on $\boldsymbol{\alpha}$
- $a, b, c$: Gompertz-Makeham distribution parameters for mortality
- $\text{S}(u) := \text{S}(u; a, b, c) = P(U \geq u)$: survival function at age $u$
- $\beta$: badger-to-badger transmission rate
- $\tau$: average latent period
- $q$: parameter to assess degree of density vs. frequency dependence
- $\nu_S, \nu_E, \nu_I$: initial probabilities of being susceptible, exposed, and infectious, respectively, at the beginning of the monitoring period
- $\xi$: Brock changepoint
- $\theta_j$: sensitivity of test $j$ during the infectious period
- $\rho_j$: scaling parameter for the test sensitivity of test $j$ during the latent period, so that $\rho_j\theta_j$ is the sensitivity of test $j$ during the latent period
- $\phi_j$: specificity of test $j$
- $\eta_\ell$: probability of capturing an alive individual at the $l$-th quarter, $\ell = 1, \ldots, 4$.
- $\boldsymbol{\Theta}$: vector including all model parameters
- $K$: rescaling constant to make the units of $\beta$ independent of $q$
- Probability distribution functions:
  - If $Z \sim \text{Exp}(a)$, $f(z) = ae^{-az}$, $a > 0$, for $z \in [0, \infty)$
  - If $Z \sim \mathcal{N}(a, b)$, $f(z) = \frac{1}{b\sqrt{2\pi}} e^{-\frac{1}{2}\left(\frac{z-a}{b}\right)^2}$, $a \in (-\infty, \infty)$, $b > 0$, for $z \in (-\infty, \infty)$
  - If $Z \sim \text{Gamma}(a, b)$, $f(z) = \frac{b^a}{\Gamma(a)} z^{a-1} e^{-bz}$, $a, b > 0$, where $\Gamma$ is the Gamma function, for $z \in (0, \infty)$
  - If $Z \sim \text{Beta}(a, b)$, $f(z) = \frac{z^{a-1}(1-z)^{b-1}}{\text{B}(a,b)}$, $a, b > 0$, where $\text{B}(a, b) = \frac{\Gamma(a)\Gamma(b)}{\Gamma(a+b)}$, for $z \in [0, 1]$

#### Coupled hidden Markov models

We use a coupled hidden Markov model (CHMM) [1] for the disease system. In this model, the unobserved state of individual $i$ at time $t$ $\left(\text{denoted by } X_t^{[i]}\right)$ depends not only on its own previous state, but also on the previous state of all other individuals. In addition, it is assumed that the observed data for individual $i$ at time $t$ only depends on the value of $X_t^{[i]}$.

The joint posterior density for the hidden states and model parameters is given by

$$\pi\left(\boldsymbol{\Theta}, \boldsymbol{X}_{1:T}^{[1:m]} \mid \boldsymbol{W}_{1:T}^{[1:m]}, \boldsymbol{o}_{1:T}^{[1:m]}, \boldsymbol{Y}_{1:T}^{[1:m]}\right) \propto \pi(\boldsymbol{\Theta}) \times$$

$$\left[\prod_{i:t_i^B<t_{0,i}} \mathrm{S}(t_{0,i}-t_i^B) P\left(X_{t_{0,i}} \mid \boldsymbol{\Theta}\right)\right] \times$$

$$\left[\prod_{i=1}^{m} \prod_{t=t_{0,i}+1}^{T_i} P\left(X_t^{[i]} \mid X_{t-1}^{[i]}, \boldsymbol{X}_{t-1}^{[-i]}, \boldsymbol{\Theta}\right)\right] \times$$

$$\left[\prod_{i=1}^{m} \prod_{t=t_{0,i}+1}^{T_i} P\Big(W_t^{[i]} \;\Big|\; X_t^{[i]}, \boldsymbol{\Theta}\Big) P\Big(\boldsymbol{Y}_t^{[i]} \;\Big|\; W_t^{[i]}, X_t^{[i]}, \boldsymbol{o}_t^{[i]}, \boldsymbol{\Theta}\Big)\right] \times$$

$$\left[\prod_{i:T_i^{\mathrm{LC}}>T_i} \prod_{t=T_i+1}^{T_i^{\mathrm{LC}}} \left(1-\psi_t^{[i]}\right)\right],$$

where the five terms are (i) the joint prior for all parameters; (ii) a survival term times a probability distribution for the states at the beginning of the monitoring period for individuals born beforehand; (iii) the probability distribution for transitions of the hidden states (these include mortality components); (iv) the probability distribution for the observation process; and (v) a survival term for individuals captured after the monitoring period. All these terms are specified below.

#### A2.1 Transition probabilities for hidden states

##### Gompertz-Makeham mortality for $\{S, E, I\} \to D$ transitions

Let $\psi_t^{[i]}$ be the probability of individual $i$ dying between $t-1$ and $t$ given it is alive at $t-1$. This will be

$$\psi_t^{[i]} := \frac{\mathrm{S}\left(\mathrm{age}_t^{[i]}-1\right) - \mathrm{S}\left(\mathrm{age}_t^{[i]}\right)}{\mathrm{S}\left(\mathrm{age}_t^{[i]}-1\right)} = 1 - \frac{\mathrm{S}\left(\mathrm{age}_t^{[i]}\right)}{\mathrm{S}\left(\mathrm{age}_t^{[i]}-1\right)}, \tag{A.1}$$

where

$$\mathrm{S}(u) := \mathrm{S}(u; a, b, c) = \exp\left\{-cu + \frac{a}{b}\left(1-e^{bu}\right)\right\}, \qquad u \in [0,\infty), \quad a, b, c > 0, \tag{A.2}$$

is the continuous-time Gompertz-Makeham survival function.

##### $S \to E$ transition probabilities given survival

We first describe the transition probability for a susceptible individual to become infected in the time period $[t-1, t)$. We assume there are two sources of infection—the background and badger-to-badger transmission. For the latter, we assume that (i) each infectious individual infects a susceptible at rate $\beta$; (ii) all infectious individuals are equally likely to infect susceptibles in the same social group; and (iii) infectious individuals are independent in $[t-1, t)$. Therefore,

$$F_{g_{i,t-1}} = \alpha_{g_i(t_i-1)} + \beta\ K^q \frac{|\mathcal{I}_{i,t_i-1}|}{|\mathcal{M}_{i,t_i-1}|^q}$$

is the force of infection acting on the susceptible individual $i$ between times $t-1$ and $t$, given it survived until time $t$. The parameter $q$ measures the strength of frequency dependence [7, 2], and $K$ is a rescaling constant that makes the units of $\beta$ independent of $q$ [7]. The value of $K$ that we used is the median social group size across all social groups and all time.

The probability of no events in $[0, t)$ in a Poisson process with rate $\Lambda$ is $\exp\{-\Lambda t\}$, motivating the choice of $\exp\{-\Lambda\}$ as the probability of avoiding infection in $[t-1, t)$ from a source with infection rate $\Lambda$. Assuming that all sources of infection are independent, the probability of avoiding infection in $[t-1, t)$ is

$$\exp\left\{-F_{g_{i,t-1}}\right\} = \exp\left\{-\alpha_{g_i(t_i-1)}\right\}\left[\prod_{j=1}^{|\mathcal{I}_{k,t_k^E-1}|} \exp\left\{-\beta\, K^q \frac{1}{|\mathcal{M}_{i,t_i-1}|^q}\right\}\right],$$

so that the probability that individual $i$ becomes infected in $[t-1, t)$, given survival until $t$, is given by

$$P\Big(X_t^{[i]} = E \,\Big|\, X_t^{[i]} \neq D,\ X_{t-1}^{[i]} = S, \boldsymbol{X}_{t-1}^{[-i]} = x_{t-1}^{[-i]}, \boldsymbol{\Theta}\Big) = 1 - \exp\left\{-F_{g_{i,t-1}}\right\}.$$

##### $E \rightarrow I$ transition probabilities given survival

We assume that an exposed individual become infectious at rate $1/\tau$, so that the discrete-time latent period has geometric distribution (given survival) with parameter $p = 1 - \exp\{-1/\tau\}$.

##### All transition probabilities

Therefore, the transition probabilities for the $i$-th individual in $[t-1, t)$ are:

$$P\Big(X_t^{[i]} = x_t^{[i]} \,\Big|\, X_{t-1}^{[i]} = x_{t-1}^{[i]}, \boldsymbol{X}_{t-1}^{[-i]} = x_{t-1}^{[-i]}, \boldsymbol{\Theta}\Big) =$$
$$\begin{cases}
\Big(1-\psi_t^{[i]}\Big)\exp\left\{-F_{g_{i,t-1}}\right\} & \text{if } x_{t-1}^{[i]} = S \text{ and } x_t^{[i]} = S \\
\Big(1-\psi_t^{[i]}\Big)\left(1-\exp\left\{-F_{g_{i,t-1}}\right\}\right) & \text{if } x_{t-1}^{[i]} = S \text{ and } x_t^{[i]} = E \\
0 & \text{if } x_{t-1}^{[i]} = S \text{ and } x_t^{[i]} = I \\
\psi_t^{[i]} & \text{if } x_{t-1}^{[i]} = S \text{ and } x_t^{[i]} = D \\
0 & \text{if } x_{t-1}^{[i]} = E \text{ and } x_t^{[i]} = S \\
\Big(1-\psi_t^{[i]}\Big)\exp\big\{-1/\tau\big\} & \text{if } x_{t-1}^{[i]} = E \text{ and } x_t^{[i]} = E \\
\Big(1-\psi_t^{[i]}\Big)\Big(1-\exp\big\{-1/\tau\big\}\Big) & \text{if } x_{t-1}^{[i]} = E \text{ and } x_t^{[i]} = I \\
\psi_t^{[i]} & \text{if } x_{t-1}^{[i]} = E \text{ and } x_t^{[i]} = D \\
0 & \text{if } x_{t-1}^{[i]} = I \text{ and } x_t^{[i]} = S \\
0 & \text{if } x_{t-1}^{[i]} = I \text{ and } x_t^{[i]} = E \\
1-\psi_t^{[i]} & \text{if } x_{t-1}^{[i]} = I \text{ and } x_t^{[i]} = I \\
\psi_t^{[i]} & \text{if } x_{t-1}^{[i]} = I \text{ and } x_t^{[i]} = D \\
0 & \text{if } x_{t-1}^{[i]} = D \text{ and } x_t^{[i]} \in \{S, E, I\} \\
1 & \text{if } x_{t-1}^{[i]} = D \text{ and } x_t^{[i]} = D
\end{cases}$$

#### A2.2 Probability distribution for the observation processes

The probability distribution of capturing badger $i$ at time $t$ is given by

$$P\Big(W_t^{[i]} = w_t^{[i]} \,\Big|\, X_t^{[i]} = x_t^{[i]}, \boldsymbol{\Theta}\Big) = \begin{cases}
\eta_{((t+4) \bmod 4)}^{w_t^{[i]}}\big(1-\eta_{((t+4) \bmod 4)}\big)^{1-w_t^{[i]}}, & \text{if } x_t^{[i]} \in \{S, E, I\} \\
1 - w_t^{[i]}, & \text{if } x_t^{[i]} = D
\end{cases}$$

where '$A$ mod $B$' refers to the modulo operation, i.e. the remainder of $A$ when divided by $B$ for integers $A$ and $B$.

The probability distribution for the test results is given by

$$P\Big(\boldsymbol{Y}_t^{[i]} = \boldsymbol{y}_t^{[i]} \;\Big|\; W_t^{[i]} = w_t^{[i]}, X_t^{[i]} = x_t^{[i]}, \boldsymbol{o}_t^{[i]}, \boldsymbol{\Theta}\Big) =$$

$$\begin{cases} \prod_{j:o_{jt}^{[i]}=1} (1-\phi_j)^{y_{jt}^{[i]}} \phi_j^{1-y_{jt}^{[i]}}, & \text{if } w_t^{[i]} = 1, \text{ tested, and } x_t^{[i]} = S \\ \prod_{j:o_{jt}^{[i]}=1} (\rho\theta_j)^{y_{jt}^{[i]}} (1-\rho\theta_j)^{1-y_{jt}^{[i]}}, & \text{if } w_t^{[i]} = 1, \text{ tested, and } x_t^{[i]} = E \\ \prod_{j:o_{jt}^{[i]}=1} \theta_j^{y_{jt}^{[i]}} (1-\theta_j)^{1-y_{jt}^{[i]}}, & \text{if } w_t^{[i]} = 1, \text{ tested, and } x_t^{[i]} = I \\ 1, & \text{if } w_t^{[i]} = 1, \text{ not tested, and } x_t^{[i]} \in \{S,E,I\} \\ 0, & \text{if } w_t^{[i]} = 1, \text{ and } x_t^{[i]} = D \\ 1, & \text{if } w_t^{[i]} = 0, \text{ and } x_t^{[i]} \in \{S,E,I,D\} \end{cases}$$

where *tested* means that the individual was tested by at least one of the tests.

Let $z_t^{[i]}$ be the Brock test result at time $t$ for individual $i$ in the model without Brock changepoint, and let $y_{1t}^{[i]}$ and $y_{2t}^{[i]}$ be the results for Brock1 and Brock2 tests, respectively, in the model with Brock changepoint. Note that

$$\begin{cases} y_{1t}^{[i]} = z_t^{[i]}, \quad o_{1t}^{[i]} = 1, \quad \text{and} \quad o_{2t}^{[i]} = 0, & \text{if } t < \xi \\ y_{2t}^{[i]} = z_t^{[i]}, \quad o_{1t}^{[i]} = 0, \quad \text{and} \quad o_{2t}^{[i]} = 1, & \text{if } t \geq \xi \end{cases} \tag{A.3}$$

Therefore, the resulting probability distribution for the observed outcomes will be

$$P\Big(W_t^{[i]} = w_t^{[i]} \;\Big|\; X_t^{[i]} = x_t^{[i]}, \boldsymbol{\Theta}\Big) \;\times\; P\Big(\boldsymbol{Y}_t^{[i]} = \boldsymbol{y}_t^{[i]} \;\Big|\; W_t^{[i]} = w_t^{[i]}, X_t^{[i]} = x_t^{[i]}, \boldsymbol{o}_t^{[i]}, \boldsymbol{\Theta}\Big) =$$

$$\begin{cases} \eta_{((t+4) \bmod 4)} \prod_{j:o_{jt}^{[i]}=1} (1-\phi_j)^{y_{jt}^{[i]}} \phi_j^{1-y_{jt}^{[i]}}, & \text{if } w_t^{[i]} = 1, \text{ tested, and } x_t^{[i]} = S \\ \eta_{((t+4) \bmod 4)} \prod_{j:o_{jt}^{[i]}=1} (\rho\theta_j)^{y_{jt}^{[i]}} (1-\rho\theta_j)^{1-y_{jt}^{[i]}}, & \text{if } w_t^{[i]} = 1, \text{ tested, and } x_t^{[i]} = E \\ \eta_{((t+4) \bmod 4)} \prod_{j:o_{jt}^{[i]}=1} \theta_j^{y_{jt}^{[i]}} (1-\theta_j)^{1-y_{jt}^{[i]}}, & \text{if } w_t^{[i]} = 1, \text{ tested, and } x_t^{[i]} = I \\ \eta_{((t+4) \bmod 4)}, & \text{if } w_t^{[i]} = 1, \text{ not tested, and } x_t^{[i]} \in \{S,E,I\} \\ 0, & \text{if } w_t^{[i]} = 1, \text{ and } x_t^{[i]} = D \\ 1 - \eta_{((t+4) \bmod 4)}, & \text{if } w_t^{[i]} = 0, \text{ and } x_t^{[i]} \in \{S,E,I\} \\ 1, & \text{if } w_t^{[i]} = 0, \text{ and } x_t^{[i]} = D \end{cases}$$

##### A2.3 Priors

We put a hierarchical shrinkage prior on the background rates of infection:

$$\begin{aligned} \alpha_g &\sim \text{Exp}(1/\lambda), \\ \lambda &\sim \text{Exp}(b_\lambda), \end{aligned} \tag{A.4}$$

$g = 1, \ldots, G$, and set $b_\lambda = 1$. On parameters $x \in \{\beta, \tau, a, b, c\}$, we used a Gamma$(a_x, b_x)$ prior, where we chose $a_\beta = a_\tau = a_a = a_b = a_c = 1$ (i.e., exponential priors), $b_\beta = b_a = b_b = b_c = 1$ and $b_\tau = 0.01$. On parameters $x \in \{q, \theta_j, \rho_j, \phi_j, \eta_\ell\}$, we used a Beta$(a_x, b_x)$ prior, where we set $a_x = b_x = 1$ in order to use a uniform prior on all these parameters. On the remaining parameters, we put priors $(\nu_S, \nu_E, \nu_I) \sim \text{Dir}(1,1,1)$ and $\xi \sim \mathcal{N}(\mu_\xi = 2000\ Q1,\ \sigma_\xi = 60 \text{ quarters})$.

### A3 Algorithm

**Algorithm** Pseudo-algorithm of iFFBS for hidden states and MCMC for model parameters.

Set $r = 1$. Draw $\boldsymbol{\Theta}$ from their initial value distributions and initialise $\boldsymbol{X}_{1:T}^{[1:m]}$ (see the rest of this section);
**for** $r = 2, \ldots, R$ **do**
  **for** $i = 1, \ldots, m$ **do**
    Draw $\boldsymbol{X}_{1:T}^{[i]} \sim P\Big(\boldsymbol{X}_{1:T}^{[i]} \mid \boldsymbol{X}_{1:T}^{[-i]}, \boldsymbol{W}_{1:T}^{[i]}, \boldsymbol{o}_{1:T}^{[i]}, \boldsymbol{Y}_{1:T}^{[i]}, \boldsymbol{\Theta}\Big)$ using iFFBS;
  **end for**
  Draw $\boldsymbol{\Theta} \sim \pi\Big(\boldsymbol{\Theta} \mid \boldsymbol{X}_{1:T}^{[1:m]}, \boldsymbol{W}_{1:T}^{[1:m]}, \boldsymbol{o}_{1:T}^{[1:m]}, \boldsymbol{Y}_{1:T}^{[1:m]}\Big)$ using appropriate MCMC (see Section A4);
**end for**

For some parameters, we used initial value distributions which were more informative than the corresponding priors: $\lambda, \beta, a, b, c, \sim$ Exp(100), $\tau \sim U(4, 16)$, $\theta_j \sim U(0.5, 1)$, $\rho_j \sim U(0.2, 0.8)$, and $\phi_j \sim U(0.7, 1)$. All the remaining parameters were initially drawn from their priors.

We also had to choose how we initialise the hidden states. For animals born during the monitoring period, we set them to be susceptible on their corresponding date of birth; for animals born beforehand, we set them to be susceptible, exposed, or infectious, with probability $\nu_S$, $\nu_E$, and $\nu_I$, respectively, at the start of the monitoring period. Each susceptible individual is set to remain susceptible in its lifetime until it is tested positive by any test, at which time we set it to be exposed. The initial latent period for each alive exposed animal is drawn from a geometric distribution with parameter $p = 1 - \exp\{-1/\tau\}$, using the initial value of $\tau$. Finally, the initial death time is set to be the first time at which the individual is no longer captured.

### A4 Parameter estimation

We used Gibbs sampling to update $\phi_j$ and $\eta_\ell$, random-walk Metropolis Hastings [6] to update $\xi$, and Hamiltonian Monte Carlo (HMC) [5] to update all the other parameters. Multidimensional adaptive random-walk Metropolis Hastings as discussed in [6] can also be used instead of HMC, but produced slower convergence and mixing.

#### A4.1 HMC updates

##### Notes on HMC for constrained parameters

This section shows how we use HMC to update a constrained parameter $x$. The idea is to use a symmetrical proposal for the transformed parameter $\tilde{x} = h(x)$ and correct the target distribution.

Suppose the target distribution is

$$\pi_x(x|D) \propto \pi_x(x) f(D|x),$$

where parameter $x$ is constrained. Typical examples are $x \in (0, 1)$, on which we put Exponential or Gamma prior, and $x \in (0, \infty)$, on which we put a Beta prior.

In the HMC, parameters are updated in terms of linear functions of derivatives:

$$x \leftarrow x + \varepsilon \frac{\partial \log \pi_x(x|D)}{\partial x}.$$

Therefore, for constrained parameters, the algorithm can easily go outside the boundaries and we would waste iterations.

We use a transformation of variable approach, so that we propose on the transformed parameter space of $\tilde{x} = h(x)$, which will be on $-\infty < \tilde{x} < \infty$. HMC steps will be of the form

$$\tilde{x} \leftarrow \tilde{x} + \varepsilon \frac{\partial \log \pi_{\tilde{x}}(\tilde{x}|D)}{\partial \tilde{x}},$$

and we need to find $\pi_{\tilde{x}}(\tilde{x}|D)$. Using transformation of variables,

$$f_{\tilde{x}}(\tilde{x}) = f_x\Big(h^{-1}(\tilde{x})\Big)|J|,$$

where $J = \dfrac{\partial x}{\partial \tilde{x}}$ is the Jacobian of the transformation.

The target distribution (i.e., the posterior of $\tilde{x}$) is therefore

$$\begin{aligned}\pi_{\tilde{x}}(\tilde{x}|D) &\propto \pi_x\left(h^{-1}(\tilde{x}) \mid D\right)\ |J| \\ &\propto \pi_x\left(h^{-1}(\tilde{x})\right) f\left(D \mid h^{-1}(\tilde{x})\right)|J|. \end{aligned} \tag{A.5}$$

The rest of this subsection will show the logarithm of the prior and Jacobian terms in Eq. (A.5), i.e. $\pi_x\left(h^{-1}(\tilde{x})\right)|J|$, as well as the derivatives with respect to the transformed parameter $\tilde{x}$, for different parameter constraints used in the paper. The likelihood terms will appear again when we discuss the posterior distributions.

**Prior and Jacobian terms for parameters on** $(0, \infty)$

For inferring a positive parameter $x > 0$, we use the logarithmic transformation $\tilde{x} = \log(x)$, and therefore the prior times Jacobian in Eq. (A.5) is given by

$$\pi_x\left(h^{-1}(\tilde{x})\right)|J| = \pi_x(e^{\tilde{x}})\ e^{\tilde{x}}.$$

Note that $\pi_x(\cdot)$ is the density function used as the prior for $x$ on the natural scale. The logarithm of the above term is therefore

$$\log\left\{\pi_x\left(h^{-1}(\tilde{x})\right)|J|\right\} = \log\left\{\pi_x(e^{\tilde{x}})\right\} + \tilde{x}.$$

Therefore, by using a Gamma prior $\pi_x(\cdot) = \text{Gamma}(a, b)$,

$$\begin{aligned}\log \pi\left\{\pi_x\left(h^{-1}(\tilde{x})\right)|J|\right\} &= \log\left\{e^{(a-1)\tilde{x}}e^{-be^{\tilde{x}}}\right\} + \tilde{x} \\ &= (a-1)\tilde{x} - be^{\tilde{x}} + \tilde{x} \\ &= a\tilde{x} - be^{\tilde{x}}, \end{aligned}$$

and

$$\frac{\partial \log\left\{\pi_x\left(h^{-1}(\tilde{x})\right)|J|\right\}}{\partial \tilde{x}} = a - be^{\tilde{x}}.$$

**Prior and Jacobian terms for parameters on** $(0, 1)$

For a parameter $x \in (0, 1)$, we use the logistic transformation $\tilde{x} = h(x) = \log\left(\dfrac{x}{1-x}\right)$, whose inverse transformation is $x = h^{-1}(\tilde{x}) = \dfrac{e^{\tilde{x}}}{1+e^{\tilde{x}}}$. In this case, the prior times Jacobian term in Eq. (A.5) is given by

$$\pi_x\left(h^{-1}(\tilde{x})\right)|J| = \pi_x\left(\frac{e^{\tilde{x}}}{1+e^{\tilde{x}}}\right)\ \frac{e^{\tilde{x}}}{(1+e^{\tilde{x}})^2}.$$

The logarithm of the above expression is thus

$$\log\left\{\pi_x\left(h^{-1}(\tilde{x})\right)|J|\right\} = \log\left\{\pi_x\left(\frac{e^{\tilde{x}}}{1+e^{\tilde{x}}}\right)\right\} + \tilde{x} - 2\log(1+e^{\tilde{x}}).$$

Therefore, by using a Beta prior $\pi_x(\cdot) = \text{Beta}(a, b)$,

$$\begin{aligned}\log\left\{\pi_x\left(h^{-1}(\tilde{x})\right)|J|\right\} &= \log\left\{\left(\frac{e^{\tilde{x}}}{1+e^{\tilde{x}}}\right)^{a-1}\left(1-\frac{e^{\tilde{x}}}{1+e^{\tilde{x}}}\right)^{b-1}\right\} + \tilde{x} - 2\log(1+e^{\tilde{x}}) \\ &= (a-1)\left[\tilde{x} - \log(1+e^{\tilde{x}})\right] + (b-1)\left[-\log(1+e^{\tilde{x}})\right] + \tilde{x} - 2\log(1+e^{\tilde{x}}) \\ &= a\tilde{x} - (a+b)\log(1+e^{\tilde{x}}) \end{aligned}$$

and

$$\frac{\partial \log \left\{ \pi_x \left( h^{-1}(\tilde{x}) \right) |J| \right\}}{\partial \tilde{x}} = a - (a+b) \left( \frac{e^{\tilde{x}}}{1+e^{\tilde{x}}} \right).$$

**Prior and Jacobian terms for the hierarchical prior on the background rates of infection**

The joint prior of $(\boldsymbol{\alpha}, \lambda)$ (see Eq. (A.4)) is given by

$$\pi(\boldsymbol{\alpha}, \lambda) \propto \pi_\lambda(\lambda; b_\lambda) \prod_{g=1}^{G} \pi_\alpha(\alpha_g | \lambda). \tag{A.6}$$

In order to improve mixing, we used a non-centred parameterisation $\alpha_g^* = h(\alpha_g) = \frac{1}{\lambda}\alpha_g$, $\forall g$. Note that

$$\begin{aligned}
\pi_{\alpha^*}(\alpha_g^* \mid \lambda) &= \pi_\alpha(h^{-1}(\alpha_g^*) \mid \lambda) \left| \frac{\partial \alpha_g}{\partial \alpha_g^*} \right| \\
&= \frac{1}{\lambda} e^{-\frac{1}{\lambda}\lambda \alpha_g^*} |\lambda| \\
&= e^{-\alpha_g^*}.
\end{aligned}$$

Therefore, $\alpha_g^* \sim \text{Exp}(1)$, and Eq. (A.6), after non-centred parameterisation, will be given by:

$$\begin{aligned}
\pi_{\boldsymbol{\alpha}^*}(\boldsymbol{\alpha}^*, \lambda) &= \pi_\lambda(\lambda; b_\lambda) \prod_{g=1}^{G} \pi_{\alpha^*}(\alpha_g^* \mid \lambda) \\
&= b_\lambda e^{-b_\lambda \lambda} \prod_{g=1}^{G} e^{-\alpha_g^*}.
\end{aligned}$$

Using log-transformations $\tilde{\alpha}_g^* = \log(\alpha_g^*)$, $\forall g$, and $\tilde{\lambda} = \log(\lambda)$, the joint prior of $(\tilde{\boldsymbol{\alpha}}_g^*, \tilde{\lambda})$ (i.e., after log-transformation) is given by

$$\begin{aligned}
\pi_{\tilde{\boldsymbol{\alpha}}^*, \tilde{\lambda}}\left(\tilde{\boldsymbol{\alpha}}_g^*, \tilde{\lambda}\right) &\propto e^{-b_\lambda e^{\tilde{\lambda}}} \left| \frac{\partial \lambda}{\partial \tilde{\lambda}} \right| \prod_{g=1}^{G} e^{-e^{\tilde{\alpha}_g^*}} \left| \frac{\partial \alpha_g^*}{\partial \tilde{\alpha}_g^*} \right| \\
&\propto e^{-b_\lambda e^{\tilde{\lambda}}} e^{\tilde{\lambda}} \prod_{g=1}^{G} e^{-e^{\tilde{\alpha}_g^*}} e^{\tilde{\alpha}_g^*},
\end{aligned}$$

whose logarithm is

$$\log\left\{\pi_{\tilde{\boldsymbol{\alpha}}^*, \tilde{\lambda}}\left(\tilde{\boldsymbol{\alpha}}_g^*, \tilde{\lambda}\right)\right\} \propto \tilde{\lambda} - b_\lambda e^{\tilde{\lambda}} + \sum_{g=1}^{G} \left(\tilde{\alpha}_g^* - e^{\tilde{\alpha}_g^*}\right).$$

The gradients used in HMC are then given by

$$\begin{aligned}
\frac{\partial \log\left\{\pi_{\tilde{\alpha}^*, \tilde{\lambda}}\left(\tilde{\alpha}_g^*, \tilde{\lambda}\right)\right\}}{\partial \tilde{\alpha}_g^*} &= 1 - e^{\tilde{\alpha}_g^*}, \quad g = 1, \ldots, G, \\
\frac{\partial \log\left\{\pi_{\tilde{\alpha}^*, \tilde{\lambda}}\left(\tilde{\alpha}_g^*, \tilde{\lambda}\right)\right\}}{\partial \tilde{\lambda}} &= 1 - b_\lambda e^{\tilde{\lambda}}.
\end{aligned}$$

##### A4.1.1 Updating $\boldsymbol{\alpha}$, $\lambda$, $\beta$, $q$, $\tau$, $a$, $b$, $c$

We used the log-transformations on $\boldsymbol{\alpha}, \lambda, \beta, \tau, a, b$, and $c$ as they are on $(0, \infty)$, and logistic transformation on $q$, which is on $(0, 1)$. The prior distributions chosen for these parameters are described in Section A2.3.

In the equations below, to simplify notation, we define

$$F_{g_{i,t-1}} := e^{\tilde{\alpha}^*_{g_i(t_i-1)}} e^{\tilde{\lambda}} + e^{\tilde{\beta}}\ K^q \frac{|\mathcal{I}_{i,t_i-1}|}{|\mathcal{M}_{i,t_i-1}|^q},$$

where $q = \dfrac{e^{\tilde{q}}}{1+e^{\tilde{q}}}$. Both $q$ and $\tilde{q}$ terms may appear in the same equation for readability.

If the individual was born before its first social group started being monitored ($t_i^B < t_{0,i}$), then the likelihood function includes the probability of having survived up to time $t_{0,i}$, i.e. $S(t_{0,i} - t_i^B)$, where $t_{0,i} - t_i^B$ is the age of individual $i$ at time $t_{0,i}$.

Time $T_i$ is the end of the monitoring period of the last social group to which individual $i$ belonged. If the last capture of individual $i$ (denoted by $T_i^{\mathrm{LC}}$) occurred after the monitoring period, this information is taken into account in the likelihood for the inference of death rates.

The joint conditional log-posterior of $(\tilde{\boldsymbol{\alpha}}, \tilde{\lambda}, \tilde{\beta}, \tilde{q}, \tilde{\tau}, \tilde{a}, \tilde{b}, \tilde{c})$ is given by:

$$\begin{aligned}
\log\Big(\pi(\tilde{\boldsymbol{\alpha}}, \tilde{\lambda}, \tilde{\beta}, \tilde{q}, \tilde{\tau}, \tilde{a}, \tilde{b}, \tilde{c} \mid \cdot\,)\Big) \propto \ & \left(\sum_{g=1}^{G}\Big(\tilde{\alpha}^*_g - e^{\tilde{\alpha}^*_g}\Big)\right) + \tilde{\lambda} - b_\lambda e^{\tilde{\lambda}} + a_\beta\tilde{\beta} - b_\beta e^{\tilde{\beta}} + \\
& a_q - (a_q + b_q)\log(1+e^{\tilde{q}}) \ + \ a_\tau\tilde{\tau} - b_\tau e^{\tilde{\tau}} \ + \ a_a\tilde{a} - b_a e^{\tilde{a}} \ + \ a_b\tilde{b} - b_b e^{\tilde{b}} + \\
& a_c\tilde{c} - b_c e^{\tilde{c}} \ + \left(\sum_{i:t_i^B < t_{0,i}} \log \mathrm{S}(t_{0,i} - t_i^B)\right) + \\
& \sum_{i=1}^{m}\sum_{t=t_{0,i}+1}^{T_i}\Bigg[ I_t^{[i]}(S,S)\Big(\log\big(1-\psi_t^{[i]}\big) - F_{g_{i,t-1}}\Big) + \\
& \quad I_t^{[i]}(S,E)\Bigg(\log\big(1-\psi_t^{[i]}\big) + \log\Big(1 - \exp\Big\{-F_{g_{i,t-1}}\Big\}\Big)\Bigg) + \\
& \quad I_t^{[i]}(E,E)\Bigg(\log\big(1-\psi_t^{[i]}\big) - 1/e^{\tilde{\tau}}\Bigg) + \\
& \quad I_t^{[i]}(E,I)\Bigg(\log\big(1-\psi_t^{[i]}\big) + \log\Big(1 - \exp\big\{-1/e^{\tilde{\tau}}\big\}\Big)\Bigg) + \\
& \quad I_t^{[i]}(1,1)\Big(\log\big(1-\psi_t^{[i]}\big)\Big) + \\
& \quad \Big(I_t^{[i]}(S,D) + I_t^{[i]}(E,D) + I_t^{[i]}(I,D)\Big)\log\Big(\psi_t^{[i]}\Big)\Bigg] + \\
& \sum_{i:T_i^{\mathrm{LC}} > T_i}\ \sum_{t=T_i+1}^{T_i^{\mathrm{LC}}} \log\Big(1-\psi_t^{[i]}\Big)
\end{aligned}$$

The gradients used in HMC are then given by

$$\begin{aligned}
\frac{\partial \log\big(\pi(\tilde{\alpha}^*_g \mid \cdot\,)\big)}{\partial \tilde{\alpha}^*_g} = \ & 1 - e^{\tilde{\alpha}^*_g} + \\
& \sum_{i=1}^{m}\sum_{t=t_{0,i}+1}^{T_i}\left[ - I_t^{[i]}(S,S)\ e^{\tilde{\alpha}^*_{g_i(t_i-1)}} e^{\tilde{\lambda}} + I_t^{[i]}(S,E)\ e^{\tilde{\alpha}^*_{g_i(t_i-1)}} e^{\tilde{\lambda}} \frac{\exp\Big\{-F_{g_{i,t-1}}\Big\}}{1-\exp\Big\{-F_{g_{i,t-1}}\Big\}}\right],
\end{aligned}$$

for whenever $g_i\,(t_i - 1) = g, \quad g = 1, \ldots, G,$

$$
\begin{aligned}
\frac{\partial \log\big(\pi(\tilde{\lambda} \mid \cdot\,)\big)}{\partial \tilde{\lambda}} &= 1 - b_\lambda e^{\tilde{\lambda}} \; + \\
&\sum_{i=1}^{m} \sum_{t=t_{0,i}+1}^{T_i} \Bigg[ - I_t^{[i]}(S,S)\; e^{\tilde{\alpha}^*_{g_i(t_i-1)}} e^{\tilde{\lambda}} + I_t^{[i]}(S,E)\; e^{\tilde{\alpha}^*_{g_i(t_i-1)}} e^{\tilde{\lambda}} \frac{\exp\Big\{-F_{g_{i,t-1}}\Big\}}{1-\exp\Big\{-F_{g_{i,t-1}}\Big\}} \Bigg], \\
\frac{\partial \log\big(\pi(\tilde{\beta} \mid \cdot\,)\big)}{\partial \tilde{\beta}} &= a_\beta - b_\beta e^{\tilde{\beta}} \; + \\
&\sum_{i=1}^{m} \sum_{t=t_{0,i}+1}^{T_i} \Bigg[ - I_t^{[i]}(S,S)\; e^{\tilde{\beta}} \left( K^q \frac{|\mathcal{I}_{i,t_i-1}|}{|\mathcal{M}_{i,t_i-1}|^q} \right) + \\
&\qquad I_t^{[i]}(S,E)\; e^{\tilde{\beta}} \left( K^q \frac{|\mathcal{I}_{i,t_i-1}|}{|\mathcal{M}_{i,t_i-1}|^q} \right) \frac{\exp\Big\{-F_{g_{i,t-1}}\Big\}}{1-\exp\Big\{-F_{g_{i,t-1}}\Big\}} \Bigg], \\
\frac{\partial \log\big(\pi(\tilde{q} \mid \cdot\,)\big)}{\partial \tilde{q}} &= a_q - (a_q + b_q) \left( \frac{e^{\tilde{q}}}{1+e^{\tilde{q}}} \right) \; + \\
&\sum_{i=1}^{m} \sum_{t=t_{0,i}+1}^{T_i} \Bigg[ I_t^{[i]}(S,S)\; e^{\tilde{\beta}} \log|\mathcal{M}_{i,t_i-1}| \left( K^q \frac{|\mathcal{I}_{i,t_i-1}|}{|\mathcal{M}_{i,t_i-1}|^q} \right) \frac{e^{\tilde{q}}}{(1+e^{\tilde{q}})^2} \; + \\
&\qquad - I_t^{[i]}(S,E)\; \frac{\exp\Big\{-F_{g_{i,t-1}}\Big\}}{1-\exp\Big\{-F_{g_{i,t-1}}\Big\}} \times \\
&\qquad\qquad e^{\tilde{\beta}} \log|\mathcal{M}_{i,t_i-1}| \left( K^q \frac{|\mathcal{I}_{i,t_i-1}|}{|\mathcal{M}_{i,t_i-1}|^q} \right) \frac{e^{\tilde{q}}}{(1+e^{\tilde{q}})^2} \Bigg], \\
\frac{\partial \log\big(\pi(\tilde{\tau} \mid \cdot\,)\big)}{\partial \tilde{\tau}} &= a_\tau - b_\tau e^{\tilde{\tau}} \; + \\
&\sum_{i=1}^{m} \sum_{t=t_{0,i}+1}^{T_i} \Bigg[ I_t^{[i]}(E,E)\; 1/e^{\tilde{\tau}} - I_t^{[i]}(E,I)\; (1/e^{\tilde{\tau}}) \frac{\exp\Big\{-1/e^{\tilde{\tau}}\Big\}}{1-\exp\Big\{-1/e^{\tilde{\tau}}\Big\}} \Bigg].
\end{aligned}
$$

In order to obtain the gradients of the survival distribution parameters, we need to know the derivatives of the survival function with respect to its parameters.

Using log-transformation on the survival distribution parameters, Eq. (A.2) becomes

$$\log \mathrm{S}(u) = -e^{\tilde{c}}u + \frac{e^{\tilde{a}}}{e^{\tilde{b}}}\big(1 - e^{e^{\tilde{b}}u}\big),$$

and therefore

$$\begin{aligned}
\frac{\partial \log \mathrm{S}(u)}{\partial \tilde{a}} &= \frac{e^{\tilde{a}}}{e^{\tilde{b}}}\big(1 - e^{e^{\tilde{b}}u}\big),\\
\frac{\partial \log \mathrm{S}(u)}{\partial \tilde{a}} &= -\frac{e^{\tilde{a}}}{e^{\tilde{b}}}\big(1 - e^{e^{\tilde{b}}u}\big) - \frac{e^{\tilde{a}}}{e^{\tilde{b}}}e^{e^{\tilde{b}}u}e^{\tilde{b}}u,\\
\frac{\partial \log \mathrm{S}(u)}{\partial \tilde{c}} &= -e^{\tilde{c}}u.
\end{aligned}$$

Let $p_t^{[i]} \coloneqq 1 - \psi_t^{[i]}$ (see Eq. (A.1)) and let $u \coloneqq \mathrm{age}_t^{[i]}$. Thus, using the log-transformation on the parameters,

$$\log p_t^{[i]} = -e^{\tilde{c}} + \frac{e^{\tilde{a}}}{e^{\tilde{b}}}\big(e^{e^{\tilde{b}}(u-1)} - e^{e^{\tilde{b}}u}\big),$$

and thus

$$\begin{aligned}
\frac{\partial \log p_t^{[i]}}{\partial \tilde{c}} &= -e^{\tilde{c}},\\
\frac{\partial \log p_t^{[i]}}{\partial \tilde{a}} &= \frac{e^{\tilde{a}}}{e^{\tilde{b}}}\big(e^{e^{\tilde{b}}(u-1)} - e^{e^{\tilde{b}}u}\big),\\
\frac{\partial \log p_t^{[i]}}{\partial \tilde{b}} &= -\frac{e^{\tilde{a}}}{e^{\tilde{b}}}\big(e^{e^{\tilde{b}}(u-1)} - e^{e^{\tilde{b}}u}\big) + e^{\tilde{a}}\Big((t-1)e^{e^{\tilde{b}}(t-1)} - te^{e^{\tilde{b}}t}\Big).
\end{aligned}$$

Finally, the derivative of $\log \psi_t^{[i]}$with respect to $\tilde{x} \in (\tilde{a}, \tilde{b}, \tilde{c})$ is given by

$$\frac{\partial \log \psi_t^{[i]}}{\partial \tilde{a}} = \frac{1}{\psi_t^{[i]}}\frac{\partial \psi_t^{[i]}}{\partial \tilde{a}} = -\frac{1}{\psi_t^{[i]}}\frac{\partial p_t^{[i]}}{\partial \tilde{a}} = -\frac{1}{\psi_t^{[i]}}p_t^{[i]}\frac{\partial \log p_t^{[i]}}{\partial \tilde{a}}.$$

Therefore,

$$\frac{\partial \log\big(\pi(\tilde{a} \mid \cdot\,)\big)}{\partial \tilde{a}} = a_a - b_a e^{\tilde{a}} + \sum_{i:t_i^B<t_{0,i}} \frac{\partial \log \mathrm{S}(t_{0,i}-t_i^B)}{\partial \tilde{a}} +$$

$$\sum_{i=1}^{m}\sum_{t=t_{0,i}+1}^{T_i}\Bigg[\Big(I_t^{[i]}(S,S)+I_t^{[i]}(S,E)+I_t^{[i]}(E,E)+I_t^{[i]}(E,I)+I_t^{[i]}(I,I)\Big)\frac{\partial \log\big(1-\psi_t^{[i]}\big)}{\partial \tilde{a}} + \Big(I_t^{[i]}(S,D)+I_t^{[i]}(E,D)+I_t^{[i]}(I,D)\Big)\frac{\partial \log \psi_t^{[i]}}{\partial \tilde{a}}\Bigg] +$$

$$\sum_{i:T_i^{\mathrm{LC}}>T_i}\ \sum_{t=T_i+1}^{T_i^{\mathrm{LC}}} \frac{\partial \log\big(1-\psi_t^{[i]}\big)}{\partial \tilde{a}},$$

$$\frac{\partial \log\big(\pi(\tilde{b} \mid \cdot\,)\big)}{\partial \tilde{b}} = a_b - b_b e^{\tilde{b}} + \sum_{i:t_i^B<t_{0,i}} \frac{\partial \log \mathrm{S}(t_{0,i}-t_i^B)}{\partial \tilde{b}} +$$

$$\sum_{i=1}^{m}\sum_{t=t_{0,i}+1}^{T_i}\Bigg[\Big(I_t^{[i]}(S,S)+I_t^{[i]}(S,E)+I_t^{[i]}(E,E)+I_t^{[i]}(E,I)+I_t^{[i]}(I,I)\Big)\frac{\partial \log\big(1-\psi_t^{[i]}\big)}{\partial \tilde{b}} + \Big(I_t^{[i]}(S,D)+I_t^{[i]}(E,D)+I_t^{[i]}(I,D)\Big)\frac{\partial \log \psi_t^{[i]}}{\partial \tilde{b}}\Bigg] +$$

$$\sum_{i:T_i^{\mathrm{LC}}>T_i}\ \sum_{t=T_i+1}^{T_i^{\mathrm{LC}}} \frac{\partial \log\big(1-\psi_t^{[i]}\big)}{\partial \tilde{b}},$$

$$\frac{\partial \log\big(\pi(\tilde{c} \mid \cdot\,)\big)}{\partial \tilde{c}} = a_c - b_c e^{\tilde{c}} + \sum_{i:t_i^B<t_{0,i}} \frac{\partial \log \mathrm{S}(t_{0,i}-t_i^B)}{\partial \tilde{c}} +$$

$$\sum_{i=1}^{m}\sum_{t=t_{0,i}+1}^{T_i}\Bigg[\Big(I_t^{[i]}(S,S)+I_t^{[i]}(S,E)+I_t^{[i]}(E,E)+I_t^{[i]}(E,I)+I_t^{[i]}(I,I)\Big)\frac{\partial \log\big(1-\psi_t^{[i]}\big)}{\partial \tilde{c}} + \Big(I_t^{[i]}(S,D)+I_t^{[i]}(E,D)+I_t^{[i]}(I,D)\Big)\frac{\partial \log \psi_t^{[i]}}{\partial \tilde{c}}\Bigg] +$$

$$\sum_{i:T_i^{\mathrm{LC}}>T_i}\ \sum_{t=T_i+1}^{T_i^{\mathrm{LC}}} \frac{\partial \log\big(1-\psi_t^{[i]}\big)}{\partial \tilde{c}}.$$

##### A4.1.2 Updating $\theta_j, \rho_j$

Using logistic transformations and putting Beta priors on $\theta_j$ and $\rho_j$ for each test $j$, the joint conditional log-posterior of $(\tilde{\theta}_j, \tilde{\rho}_j)$ is given by

$$\begin{aligned}
\log\Big(\pi(\tilde{\theta}_j, \tilde{\rho}_j | \cdots)\Big) &\propto a_{\tilde{\theta}}\tilde{\theta}_j - (a_{\tilde{\theta}} + b_{\tilde{\theta}})\log(1+e^{\tilde{\theta}_j}) \; + \; a_{\tilde{\rho}}\tilde{\rho}_j - (a_{\tilde{\rho}} + b_{\tilde{\rho}})\log(1+e^{\tilde{\rho}_j}) \; + \\
&\sum_{i=1}^{m}\sum_{o_{jt}^{[i]}=1}\Bigg\{\Bigg[y_{jt}^{[i]}\log\left(\frac{e^{\tilde{\theta}_j}}{1+e^{\tilde{\theta}_j}}\right) + (1-y_{jt}^{[i]})\log\left(1-\frac{e^{\tilde{\theta}_j}}{1+e^{\tilde{\theta}_j}}\right)\Bigg]\mathbb{1}_{\{X_t^{[i]}=I\}} + \\
&\Bigg[y_{jt}^{[i]}\left(\log\left(\frac{e^{\tilde{\theta}_j}}{1+e^{\tilde{\theta}_j}}\right) + \log\left(\frac{e^{\tilde{\rho}_j}}{1+e^{\tilde{\rho}_j}}\right)\right) + \\
&(1-y_{jt}^{[i]})\log\left(1-\frac{e^{\tilde{\theta}_j}}{1+e^{\tilde{\theta}_j}}\frac{e^{\tilde{\rho}_j}}{1+e^{\tilde{\rho}_j}}\right)\Bigg]\mathbb{1}_{\{X_t^{[i]}=E\}}\Bigg\} \\
&\propto a_{\tilde{\theta}}\tilde{\theta}_j - (a_{\tilde{\theta}} + b_{\tilde{\theta}})\log(1+e^{\tilde{\theta}_j}) \; + \; a_{\tilde{\rho}}\tilde{\rho}_j - (a_{\tilde{\rho}} + b_{\tilde{\rho}})\log(1+e^{\tilde{\rho}_j}) \; + \\
&\sum_{i=1}^{m}\sum_{o_{jt}^{[i]}=1}\Bigg\{\Big[y_{jt}^{[i]}\Big(\tilde{\theta}_j - \log(1+e^{\tilde{\theta}_j})\Big) - (1-y_{jt}^{[i]})\log(1+e^{\tilde{\theta}_j})\Big]\mathbb{1}_{\{X_t^{[i]}=I\}} + \\
&\Bigg[y_{jt}^{[i]}\Big(\tilde{\theta}_j - \log(1+e^{\tilde{\theta}_j}) + \tilde{\rho}_j - \log(1+e^{\tilde{\rho}_j})\Big) + \\
&(1-y_{jt}^{[i]})\Big(\log(1+e^{\tilde{\theta}_j}+e^{\tilde{\rho}_j}) - \log(1+e^{\tilde{\theta}_j}) - \log(1+e^{\tilde{\rho}_j})\Big)\Bigg]\mathbb{1}_{\{X_t^{[i]}=E\}}\Bigg\}.
\end{aligned}$$

The gradients used in HMC are therefore given by

$$\begin{aligned}
\frac{\partial\log\Big(\pi(\tilde{\theta}_j|\cdots)\Big)}{\partial\tilde{\theta}_j} &= a_\theta - (a_\theta + b_\theta)\frac{e^{\tilde{\theta}_j}}{1+e^{\tilde{\theta}_j}} \; + \\
&\sum_{i=1}^{m}\sum_{o_{jt}^{[i]}=1}\Bigg\{\Bigg(y_{jt}^{[i]}\left(1-\frac{e^{\tilde{\theta}_j}}{1+e^{\tilde{\theta}_j}}\right) - (1-y_{jt}^{[i]})\frac{e^{\tilde{\theta}_j}}{1+e^{\tilde{\theta}_j}}\Bigg)\mathbb{1}_{\{X_t^{[i]}=I\}} \; + \\
&\Bigg(y_{jt}^{[i]}\left(1-\frac{e^{\tilde{\theta}_j}}{1+e^{\tilde{\theta}_j}}\right) + (1-y_{jt}^{[i]})\left(\frac{e^{\tilde{\theta}_j}}{1+e^{\tilde{\theta}_j}+e^{\tilde{\rho}_j}} - \frac{e^{\tilde{\theta}_j}}{1+e^{\tilde{\theta}_j}}\right)\Bigg)\mathbb{1}_{\{X_t^{[i]}=E\}}\Bigg\},
\end{aligned}$$

and

$$\begin{aligned}
\frac{\partial\log\Big(\pi(\tilde{\rho}_j|\cdots)\Big)}{\partial\tilde{\rho}_j} &= a_\rho - (a_\rho + b_\rho)\frac{e^{\tilde{\rho}_j}}{1+e^{\tilde{\rho}_j}} \; + \\
&\sum_{i=1}^{m}\sum_{o_{jt}^{[i]}=1}\Bigg\{\Bigg(y_{jt}^{[i]}\left(1-\frac{e^{\tilde{\rho}_j}}{1+e^{\tilde{\rho}_j}}\right) + (1-y_{jt}^{[i]})\left(\frac{e^{\tilde{\rho}_j}}{1+e^{\tilde{\theta}_j}+e^{\tilde{\rho}_j}} - \frac{e^{\tilde{\rho}_j}}{1+e^{\tilde{\rho}_j}}\right)\Bigg)\mathbb{1}_{\{X_t^{[i]}=E\}}\Bigg\}.
\end{aligned}$$

#### A4.2 Gibbs sampling updates

In this section, we show the conditional distributions used in Gibbs sampling updates for the remaining parameters. Priors choices are described in Section A2.3.

###### A4.2.1 Updating $\phi_j$

The conditional posterior distribution of $\phi_j$ is given by

$$\pi\big(\phi_j \bigm| \cdots\big) \propto \phi_j^{a_{\phi_j}-1}(1-\phi_j)^{b_{\phi_j}-1}\prod_{i=1}^{m}\prod_{\substack{X_t^{[i]}=S;\\ o_{jt}^{[i]}=1}}\Big[(1-\phi_j)^{y_{jt}^{[i]}}\phi_j^{1-y_{jt}^{[i]}}\Big].$$

Therefore, in the Gibbs step,

$$\phi_j \sim \text{Beta}\Big(a_{\phi_j}+\sum_{i=1}^{m}\sum_{\substack{X_t^{[i]}=S;\\ o_{jt}^{[i]}=1}}\big(1-y_{jt}^{[i]}\big),\ \ b_{\phi_j}+\sum_{i=1}^{m}\sum_{\substack{X_t^{[i]}=S;\\ o_{jt}^{[i]}=1}} y_{jt}^{[i]}\Big).$$

###### A4.2.2 Updating $\eta_\ell$

For conditional posterior distribution for the quarterly capture probability $\eta_\ell,\ \ell=1,\ldots,4$, is given by

$$\pi\big(\eta_\ell \bigm| \cdots\big) \propto\ \eta_\ell^{a_{\eta_\ell}-1}(1-\eta_\ell)^{b_{\eta_\ell}-1}\prod_{i=1}^{m}\prod_{\substack{t=1;\\ X_t^{[i]}\in\{S,E,I\}}}^{T} \mathbb{1}_{\{4|(t-\ell+4)\}}\Big[\eta_\ell^{w_t^{[i]}}(1-\eta_\ell)^{1-w_t^{[i]}}\Big]$$

$$\propto \eta_\ell^{a_{\eta_\ell}-1+\sum_{i=1}^{m}\sum_{t=1;X_t^{[i]}\in\{S,E,I\}}^{T}\mathbb{1}_{\{4|(t-\ell+4)\}}w_t^{[i]}}\ (1-\eta_\ell)^{b_{\eta_\ell}-1+\sum_{t=1;X_t^{[i]}\in\{S,E,I\}}^{m}\sum_{t=1}^{T}\mathbb{1}_{\{4|(t-\ell+4)\}}(1-w_t^{[i]})},$$

where the indicator function $\mathbb{1}_{\{4|(t-\ell+4)\}}$ equals 1 if $t-\ell+4$ is a multiple of 4, and 0 otherwise. Therefore, the Gibbs step consists of drawing from

$$\eta_\ell \sim \text{Beta}\Big(a_{\eta_\ell}+\sum_{i=1}^{m}\sum_{\substack{t=1;\\ X_t^{[i]}\in\{S,E,I\}}}^{T}\mathbb{1}_{\{4|(t-\ell+4)\}}w_t^{[i]},\ b_{\eta_\ell}+\sum_{i=1}^{m}\sum_{\substack{t=1;\\ X_t^{[i]}\in\{S,E,I\}}}^{T}\mathbb{1}_{\{4|(t-\ell+4)\}}(1-w_t^{[i]})\Big).$$

###### A4.2.3 Updating $\nu_S, \nu_E, \nu_I$

The joint conditional posterior distribution of $(\nu_S, \nu_E, \nu_I)$, by using a Dirichlet prior Dir(1, 1, 1), is

$$\pi\big(\nu_S,\nu_E,\nu_I \bigm| \cdots\big) \propto \prod_{i:t_i^B<t_{0,i}}\Big[\nu_S^{\mathbb{1}_{\{X_{t_{0,i}}^{[i]}=S\}}}\nu_E^{\mathbb{1}_{\{X_{t_{0,i}}^{[i]}=E\}}}\nu_I^{\mathbb{1}_{\{X_{t_{0,i}}^{[i]}=I\}}}\Big].$$

Note that we use only information from the individuals which were born before the monitoring period, i.e. $t_i^B < t_{0,i}$. Thus, values for $(\nu_S, \nu_E, \nu_I)$ are sampling from

$$(\nu_S,\nu_E,\nu_I) \sim \text{Dir}\Big(1+\sum_{i:t_i^B<t_{0,i}}\mathbb{1}_{\{X_{t_{0,i}}^{[i]}=S\}},\ 1+\sum_{i:t_i^B<t_{0,i}}\mathbb{1}_{\{X_{t_{0,i}}^{[i]}=E\}},\ 1+\sum_{i:t_i^B<t_{0,i}}\mathbb{1}_{\{X_{t_{0,i}}^{[i]}=I\}}\Big).$$

##### A4.3 Metropolis-Hastings update for $\xi$

Brock changepoint $\xi$ is the only parameter updated by Metropolis-Hastings. The conditional log-posterior of $\xi$ is given by

$$\log\left(\pi(\xi|\cdots)\right) \propto -\frac{1}{2\sigma_\xi^2}(\xi-\mu_\xi)^2 + \sum_{i=1}^{m}\sum_{o_{jt}^{[i]}=1}\Bigg\{\left[y_{jt}^{[i]}\log(\theta_j)+(1-y_{jt}^{[i]})\log(1-\theta_j)\right]\mathbb{1}_{\{X_t^{[i]}=I\}} + \left[y_{jt}^{[i]}\log(\rho_j\theta_j)+(1-y_{jt}^{[i]})\log(1-\rho_j\theta_j)\right]\mathbb{1}_{\{X_t^{[i]}=E\}} + \left[y_{jt}^{[i]}\log(1-\phi_j)+(1-y_{jt}^{[i]})\log(\phi_j)\right]\mathbb{1}_{\{X_t^{[i]}=S\}}\Bigg\}$$

where $o_{jt}^{[i]}$ and $y_{jt}^{[i]}$, for tests $j \in \{1,2\}$, depend on $\xi$ (see Eq. (A.3)) in the model with Brock changepoint.

#### A5 Individual effective reproduction number

[8] derive a likelihood-based estimator for the individual effective reproduction number, $R_i$, for an individual $i$ as:

$$R_i = \sum_{k|t_k^E > t_i^E} p_{(k,i)}. \tag{A.7}$$

Here $p_{(k,i)}$ is the relative likelihood of individual $k$ being infected by individual $i$, and hence $R_i$ is the sum of these relative likelihoods over all possible future infections $\left\{k; t_k^E > t_i^E\right\}$. [8] derive an estimator for $p_{(k,i)}$ based on the generation-time distribution $w(\cdot)$, such that:

$$p_{(k,i)} = \frac{w\left(t_k^E - t_i^E\right)}{\sum_{j|t_k^E > t_j^E} w\left(t_k^E - t_j^E\right)}.$$

Ordinarily, since the specific who-infected-whom infection network is unknown it is necessary to integrate over all possible infection networks to generate an estimate of $R_i$, which is computationally intractable unless very few individuals are infected. However, under the assumption that the prior probability of any given infection network being true is constant, [8] show that (A.7) is equivalent to integrating over all possible networks, and is substantially more efficient.

Since our model is not defined in terms of the generation-time distribution, we instead derive a variant of this estimator that utilises the exact likelihoods that we have in our model. Furthermore, to use the [8] estimator directly also requires knowledge of the infection times, which are unknown. [8] used the difference between reported dates of symptom onset as a proxy for $t_k^E - t_i^E$. Our approach does not need any proxy, as the iFFBS algorithm generates random samples for the unobserved event times at each iteration of the MCMC algorithm, and we can condition on these directly to generate random samples from the posterior distribution of $R_i$. Taken over the whole MCMC chain, these can be used to estimate the posterior distribution for $R_i$ which numerically integrates over the relevant hidden states. In addition, we adjust for the fact that we have additional sources of infectious pressure through the background rates-of-infection.

##### A5.1 Discrete event times

At a given iteration of the MCMC, we have imputed infection times, $t_k^E$, for $N^E$ infected individuals ($k = 1, \ldots, N^E$). In practice $N^E$ may change between iterations, since it depends on the infection states of individuals, which are unknown and thus imputed using the iFFBS algorithm. For brevity, we ignore any dependence on the specific MCMC iteration in the discussion below.

Hence, at a given iteration we have a set of simulated discrete infection times, $t_k^E$. From [8] we can define:

$$R_i = \sum_{k: t_k^E > t_i^E} p_{(k,i)}, \tag{A.8}$$

where $p_{(k,i)}$ is the relative likelihood that individual $i$ infects individual $k$. Using the same assumptions as [8], and assuming no background rate-of-infection terms, this reduces to:

$$p_{(k,i)} = \frac{P_{ki}\left(t_k^E - 1 < T_k^E < t_k^E \mid T_k^E > t_k^E - 1\right) \frac{1}{|\mathcal{I}_{k,t_k^E-1}|} \prod_{t=t_k^B+1}^{t_k^E-1} P_{ki}\left(T_k^E > t \mid T_k^E > t-1\right)}{\sum_{j\in\mathcal{I}_{k,t_k^E-1}} P_{kj}\left(t_k^E - 1 < T_k^E < t_k^E \mid T_k^E > t_k^E - 1\right) \frac{1}{|\mathcal{I}_{k,t_k^E-1}|} \prod_{t=t_k^B+1}^{t_k^E-1} P_{kj}\left(T_k^E > t \mid T_k^E > t-1\right)} \tag{A.9}$$

which simplifies to:

$$p_{(k,i)} = \begin{cases} \dfrac{1}{|\mathcal{I}_{k,t_k^E-1}|} & \text{if } i \in \mathcal{I}_{k,t_k^E-1}, \\ 0 & \text{otherwise.} \end{cases}$$

Here, $P_{ki}(\cdot)$ correspond to probability statements regarding infection events for individual $k$ due to the infectious pressure generated by individual $i$.

In (A.9), $\dfrac{1}{|\mathcal{I}_{k,t_k^E-1}|}$ refers to the probability that individual $k$ was infected in $\left[t_k^E - 1, t_k^E\right)$ by one of the individuals which were infectious at $t_k^E - 1$ in the same social group, given that individual $k$ was susceptible at $t_k^E - 1$. This comes from the fact that each individual $j \in \mathcal{I}_{k,t_k^E-1}$ is assumed to have the same probability of infecting $k$.

##### Adjusting for background rates-of-infection

For our model we also need to adjust the estimator for $R_i$ to account for infectious pressure due to the background rate-of-infection terms. Similarly to Eq. (A.9) we can consider:

$$p_{(k,i)} \propto P_{ki}\left(t_k^E - 1 < T_k^E < t_k^E \mid T_k^E > t_k^E - 1\right) \frac{\beta K^q / |\mathcal{M}_{k,t_k^E-1}|^q}{\alpha_{g_k(t_k^E-1)} + |\mathcal{I}_{k,t_k^E-1}| \beta K^q / |\mathcal{M}_{k,t_k^E-1}|^q} \prod_{t=t_k^B+1}^{t_k^E-1} P_{ki}\left(T_k^E > t \mid T_k^E > t-1\right),$$

where the constant-of-proportionality is given by:

$$P_{k\alpha}\left(t_k^E - 1 < T_k^E < t_k^E \mid T_k^E > t_k^E - 1\right) \frac{\alpha_{g_k(t_k^E-1)}}{\alpha_{g_k(t_k^E-1)} + |\mathcal{I}_{k,t_k^E-1}| \beta K^q / |\mathcal{M}_{k,t_k^E-1}|^q} \prod_{t=t_k^B+1}^{t_k^E-1} P_{k\alpha}\left(T_k^E > t \mid T_k^E > t-1\right) +$$

$$\sum_{j\in\mathcal{I}_{k,t_k^E-1}} \left[P_{ki}\left(t_k^E - 1 < T_k^E < t_k^E \mid T_k^E > t_k^E - 1\right) \frac{\beta K^q / |\mathcal{M}_{k,t_k^E-1}|^q}{\alpha_{g_k(t_k^E-1)} + |\mathcal{I}_{k,t_k^E-1}| \beta K^q / |\mathcal{M}_{k,t_k^E-1}|^q} \prod_{t=t_k^B+1}^{t_k^E-1} P_{ki}\left(T_k^E > t \mid T_k^E > t-1\right)\right].$$

Therefore,

$$p_{(k,i)} = \begin{cases} \dfrac{\beta K^q / |\mathcal{M}_{k,t_k^E-1}|^q}{\alpha_{g_k(t_k^E-1)} + |\mathcal{I}_{k,t_k^E-1}| \beta K^q / |\mathcal{M}_{k,t_k^E-1}|^q} & \text{if } i \in \mathcal{I}_{k,t_k^E-1}, \\ 0 & \text{otherwise.} \end{cases}$$

##### Illustration

For simplicity, assume that all the $m$ individuals remain alive and belong to the same social group $g$. Let $t_i^E$ and $t_i^I$ be the times at which individual $i$ becomes exposed and infectious, respectively.

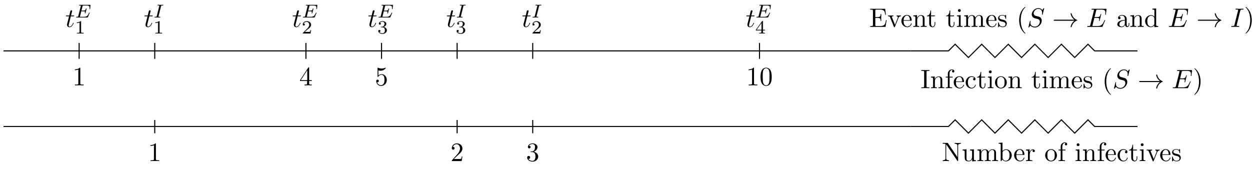

Let $p_{(k,i)}$ be the probability that badger $i$ infects badger $k$ at each new infection time $t_k^E$, and let $p_{(k,\alpha_g)}$ be the probability that badger $k$ is infected via the background rate-of-transmission term. Below we illustrate how $R_i$ is obtained for the individuals that became infectious ($i = 1, 2, 3$).

| Newly infected badger ($k$) | 1 | 2 | 3 | 4 |
|---|---|---|---|---|
| Infection time ($t_k^E$) | 1 | 4 | 5 | 10 |
| Total rate ($r_{t-1}$) | $\alpha_g$ | $\alpha_g + \beta\frac{K^q}{m^q}$ | $\alpha_g + \beta\frac{K^q}{m^q}$ | $\alpha_g + 3\beta\frac{K^q}{m^q}$ |
| $p_{(k,1)}$ | 0 | $\beta\frac{K^q}{m^q}/r_3$ | $\beta\frac{K^q}{m^q}/r_4$ | $\beta\frac{K^q}{m^q}/r_9$ |
| $p_{(k,2)}$ | 0 | 0 | 0 | $\beta\frac{K^q}{m^q}/r_9$ |
| $p_{(k,3)}$ | 0 | 0 | 0 | $\beta\frac{K^q}{m^q}/r_9$ |
| $p_{(k,\alpha_g)}$ | 1 | $\alpha_g/r_3$ | $\alpha_g/r_4$ | $\alpha_g/r_9$ |

We define the individual reproduction number, $R_i$ from (A.8) as:

$$R_1 = \beta\frac{K^q}{m^q}\left(\frac{1}{r_3} + \frac{1}{r_4} + \frac{1}{r_9}\right),$$

$$R_2 = \beta\frac{K^q}{m^q}\left(\frac{1}{r_9}\right),$$

$$R_3 = \beta\frac{K^q}{m^q}\left(\frac{1}{r_9}\right).$$

The estimate of the posterior distribution for each $R_i$ is obtained from the values obtained for each iteration of the MCMC. The population reproduction number is defined as the arithmetic mean of $R_i$ over all badgers.

We note that it is in theory possible to derive an effective $R_{\alpha_g}$ here for the background rate-of-infection terms, but since the background rates of infection are constant in this model, then $R_{\alpha_g}$ is not constrained by any kind of infectious period, and so from (A.8) we can see that $R_{\alpha_g}$ would simply increase as the total number of infected badgers increases, so it does not make much biological sense to do this here.

#### A5.2 Population-level effective reproduction numbers

There are various ways in which population-level effective reproduction numbers can be defined, see for example [8, 4, 3]. Here we do something more akin to the approach of [8], in which we retrospectively generate the expected number of cases infected by an animal across its lifetime. To obtain a population-level estimate, we simply average across individuals, giving:

$$R_t = \frac{1}{N_t^D}\sum_{i=1}^{N_t^D} R_i,$$

where $R_i$ is defined in (A.8) and $N_t^D$ is the number of badgers that have died up to time $t$. Hence $R_t$ is *retrospective*, in the sense that it is an estimate of the expected number of secondary infections that were caused by a random infected individual across its whole lifetime. We restrict to only looking at individuals who have died in order to keep within the paradigm of trying to understand what has happened in the population so far, rather than what will happen going forwards. For the latter we could perform forwards-simulations from the model under some assumptions about future movements of animals, that would have to be determined.

#### A6 Relative rates of infection

For each iteration of the MCMC, the relative contribution of direct badger-to-badger transmission on any infected badger $k$ at its infection time $t_k^E$ can be calculated as:

$$\frac{\beta K^q |\mathcal{I}_{k,t_k^E-1}| / |\mathcal{M}_{k,t_k^E-1}|^q}{\alpha_{g_k\left(t_k^E-1\right)} + \beta K^q |\mathcal{I}_{k,t_k^E-1}| / |\mathcal{M}_{k,t_k^E-1}|^q}.$$

These can then be aggregated at any spatio-temporal resolution, and repeating this process for different MCMC samples allows for a predictive distribution to be built empirically.

### A7 Additional results

#### A7.1 Mixing

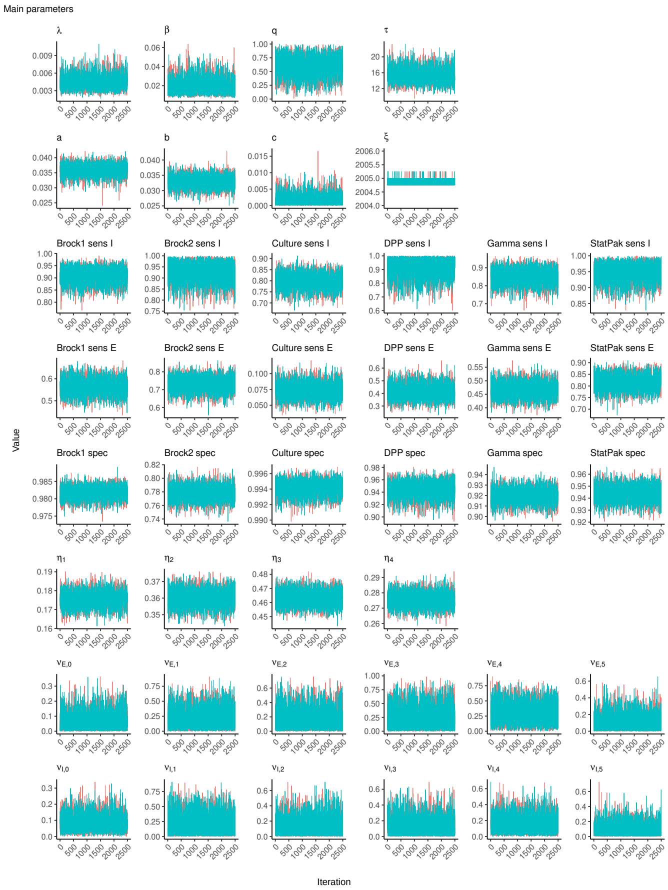

Figure A1: Traceplots of all parameters (except background rates of infection).

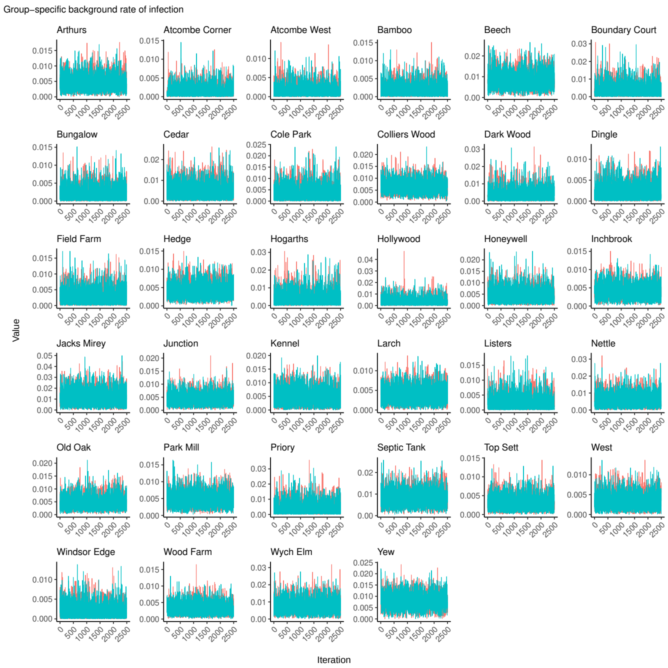

Figure A2: Traceplots of group-specific background rates of infection.

#### A7.2 Additional model outputs

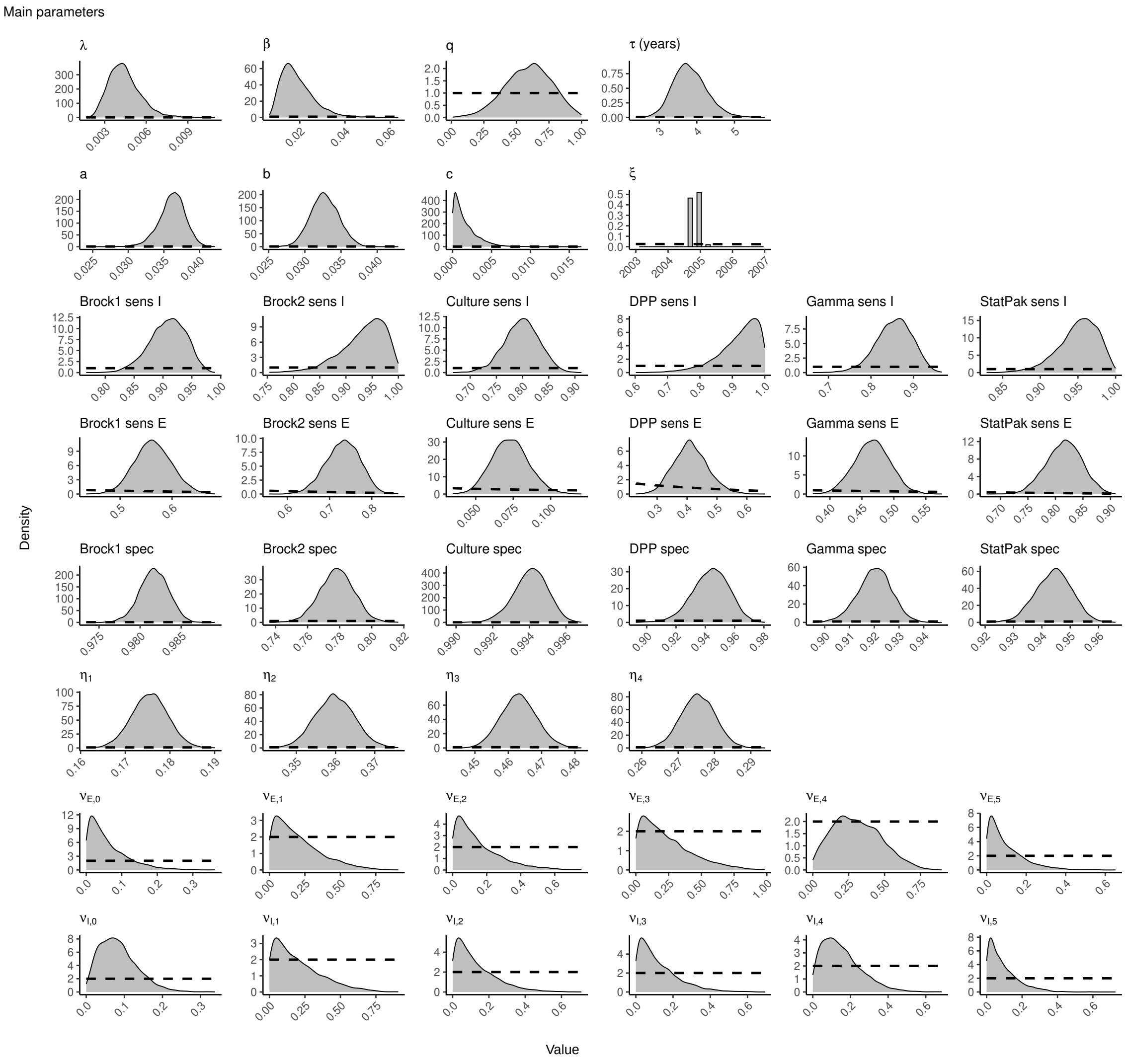

Figure A3: Posterior densities for all parameters except the background rates of infection. Dashed lines represent the prior densities.

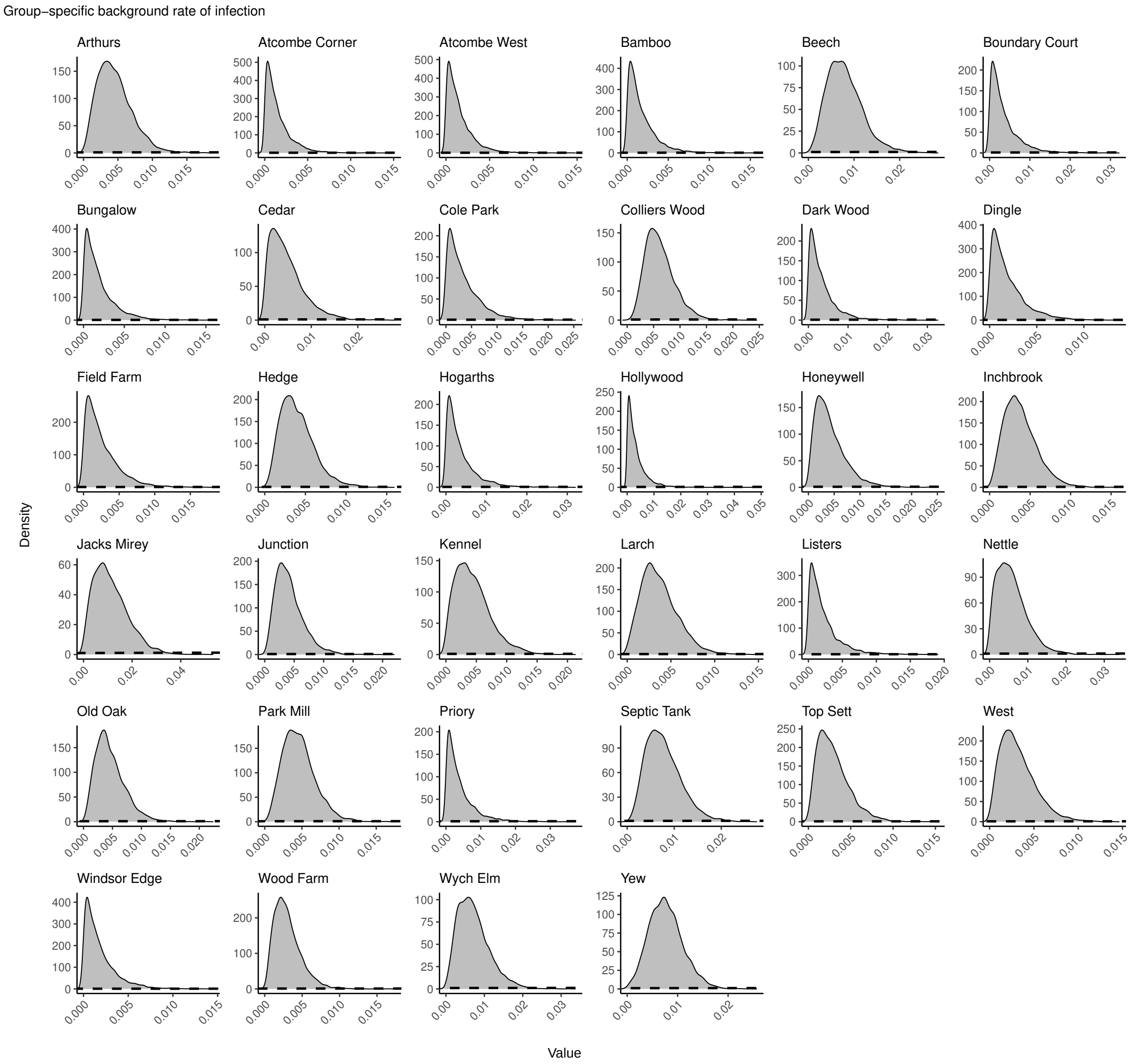

Figure A4: Posterior densities of the background rates of infection. Dashed lines represent the prior densities.

| Group | Mean | 95% CI |
|---|---|---|
| Arthurs | 0.0045 | (0.00088, 0.01) |
| Atcombe Corner | 0.0015 | (0.000036, 0.0054) |
| Atcombe West | 0.0015 | (0.000037, 0.0054) |
| Bamboo | 0.0017 | (0.000049, 0.0062) |
| Beech | 0.008 | (0.002, 0.017) |
| Boundary Court | 0.0034 | (0.000082, 0.013) |
| Bungalow | 0.0019 | (0.000046, 0.0068) |
| Cedar | 0.0047 | (0.00027, 0.014) |
| Cole Park | 0.0034 | (0.000084, 0.012) |
| Colliers Wood | 0.0062 | (0.002, 0.013) |
| Dark Wood | 0.0033 | (0.000085, 0.012) |
| Dingle | 0.0019 | (0.000051, 0.0068) |
| Field Farm | 0.0025 | (0.000077, 0.0086) |
| Hedge | 0.004 | (0.001, 0.0089) |
| Hogarths | 0.0036 | (0.000081, 0.014) |
| Hollywood | 0.0032 | (0.000063, 0.013) |
| Honeywell | 0.0042 | (0.00049, 0.012) |
| Inchbrook | 0.0038 | (0.00081, 0.0085) |
| Jacks Mirey | 0.011 | (0.0012, 0.028) |
| Junction | 0.0042 | (0.00087, 0.0098) |
| Kennel | 0.0044 | (0.00036, 0.012) |
| Larch | 0.0037 | (0.00055, 0.0084) |
| Listers | 0.0022 | (0.000061, 0.0084) |
| Nettle | 0.0058 | (0.00039, 0.015) |
| Old Oak | 0.0046 | (0.001, 0.011) |
| Park Mill | 0.0046 | (0.0012, 0.0095) |
| Priory | 0.0039 | (0.00011, 0.015) |
| Septic Tank | 0.0077 | (0.0021, 0.016) |
| Top Sett | 0.0029 | (0.00043, 0.0074) |
| West | 0.0032 | (0.00046, 0.008) |
| Windsor Edge | 0.0018 | (0.000048, 0.0066) |
| Wood Farm | 0.003 | (0.00058, 0.0073) |
| Wych Elm | 0.0072 | (0.0012, 0.017) |
| Yew | 0.0077 | (0.0019, 0.015) |

Table A1: Posterior means and 95% credible intervals for group-specific background rate-of-infection terms. Shown to 2 significant figures.

#### A7.3 Social group model fits

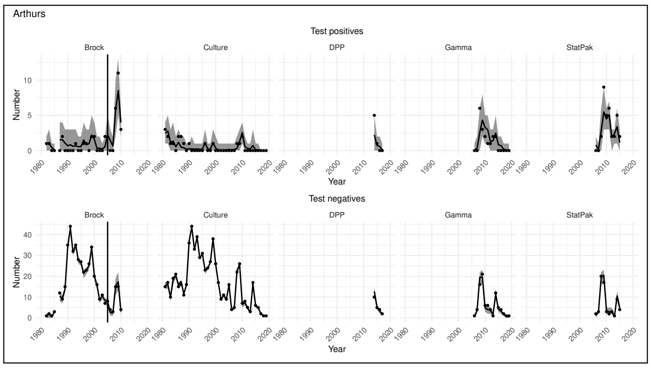

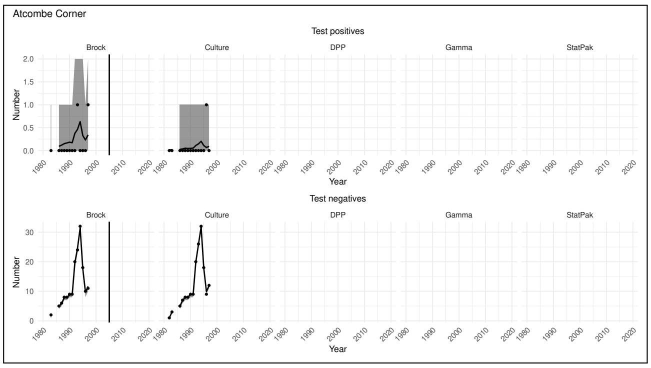

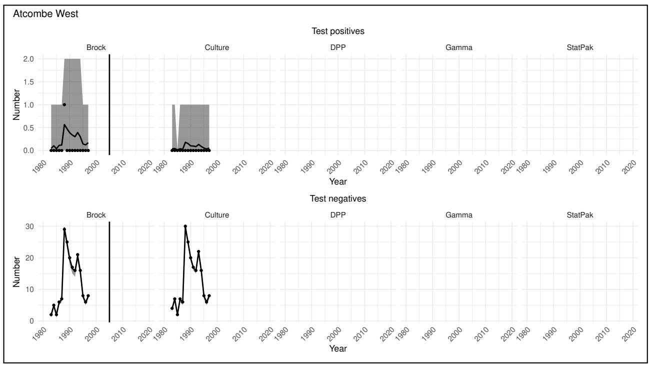

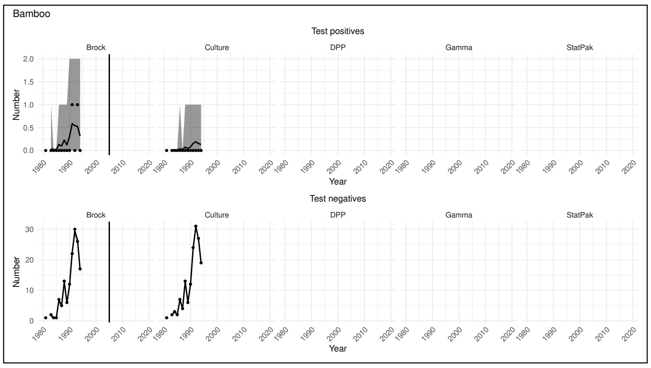

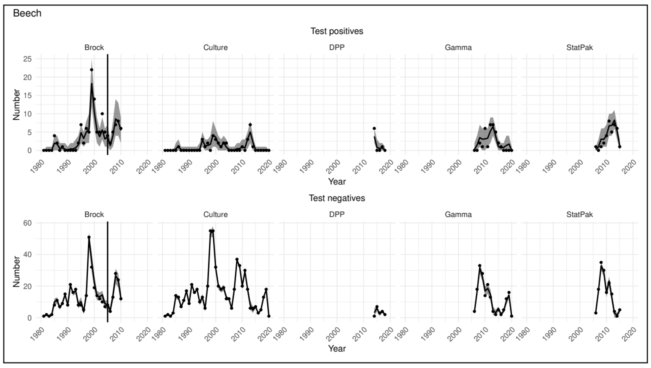

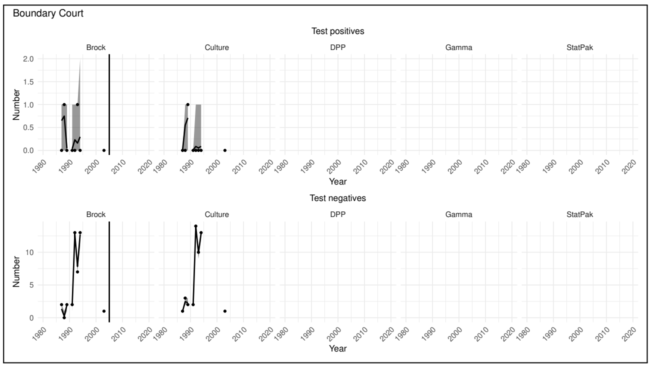

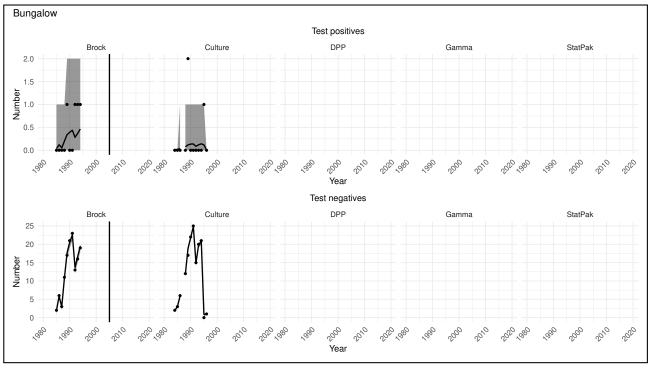

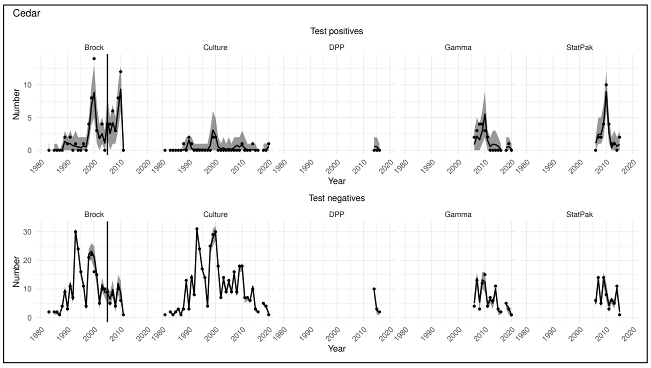

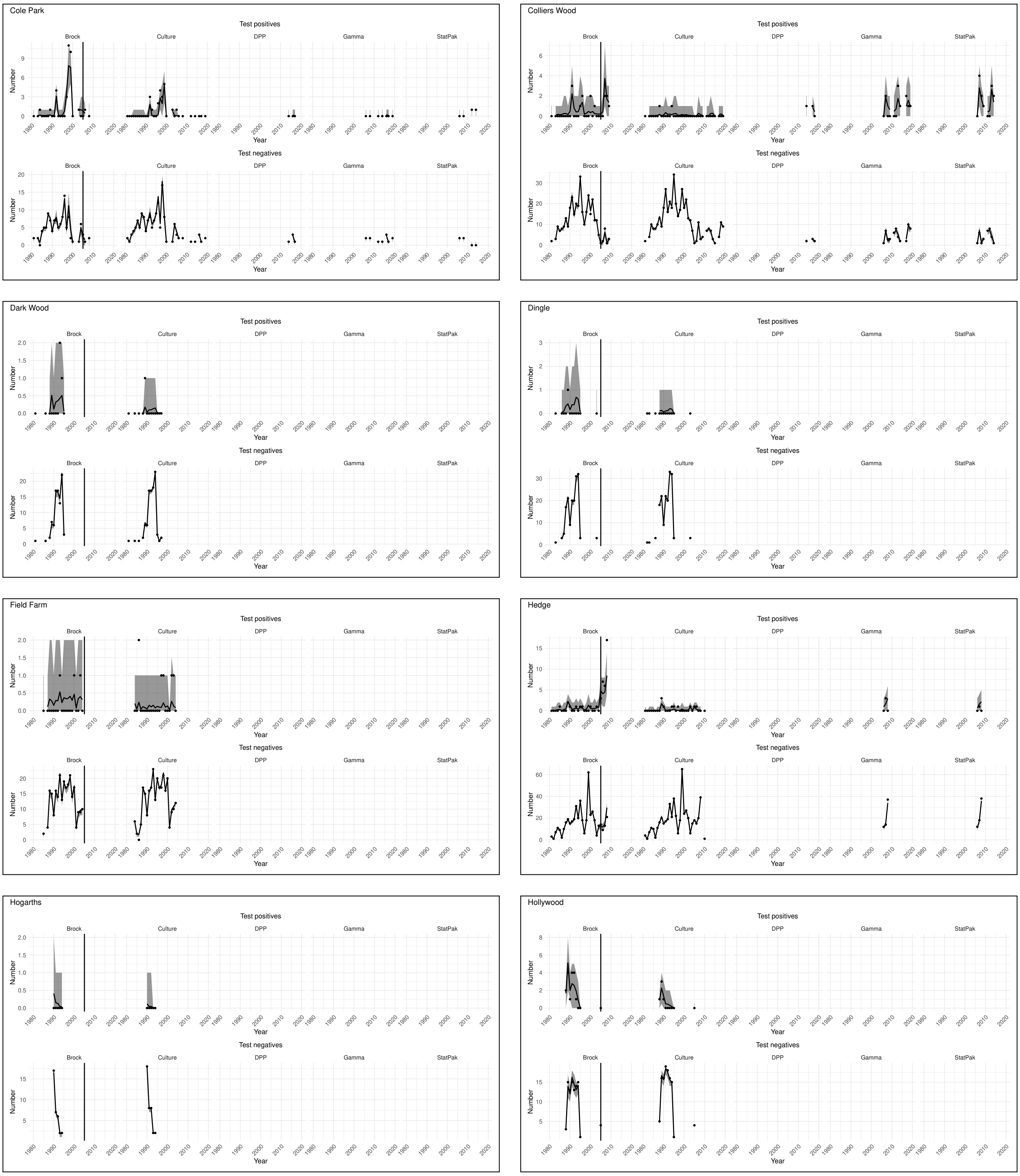
Cole Park
Colliers Wood
Dark Wood
Dingle
Field Farm
Hedge
Hogarths
Hollywood
Test positives
Test negatives
Brock
Culture
DPP
Gamma
StatPak
Number
Year

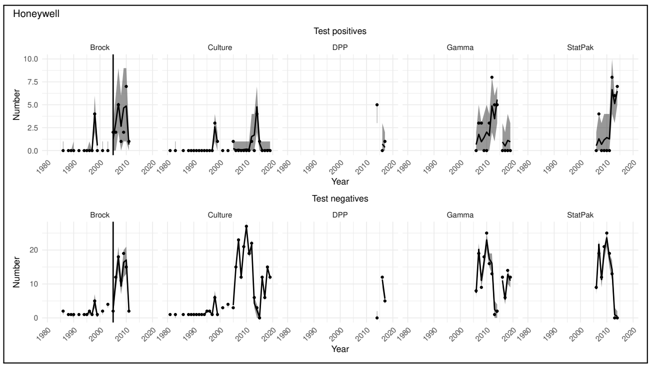
Honeywell
Test positives
Brock
Culture
DPP
Gamma
StatPak
Number
Year
Test negatives

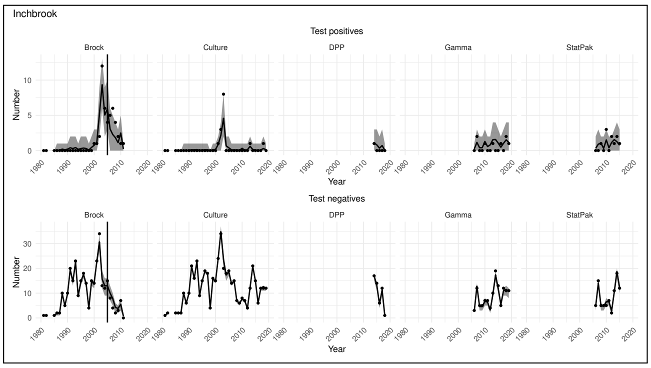
Inchbrook
Test positives
Brock
Culture
DPP
Gamma
StatPak
Number
Year
Test negatives

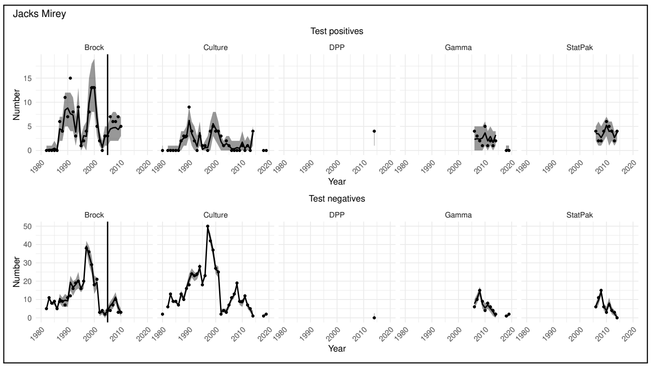
Jacks Mirey
Test positives
Brock
Culture
DPP
Gamma
StatPak
Number
Year
Test negatives

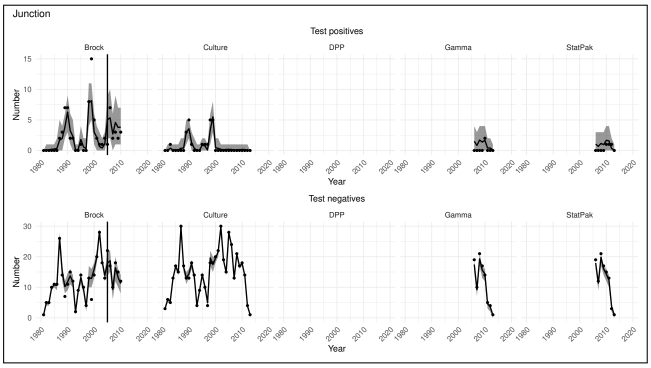
Junction
Test positives
Brock
Culture
DPP
Gamma
StatPak
Number
Year
Test negatives

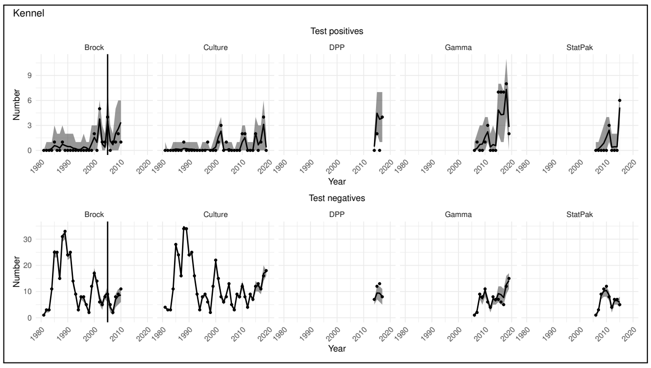
Kennel
Test positives
Brock
Culture
DPP
Gamma
StatPak
Number
Year
Test negatives

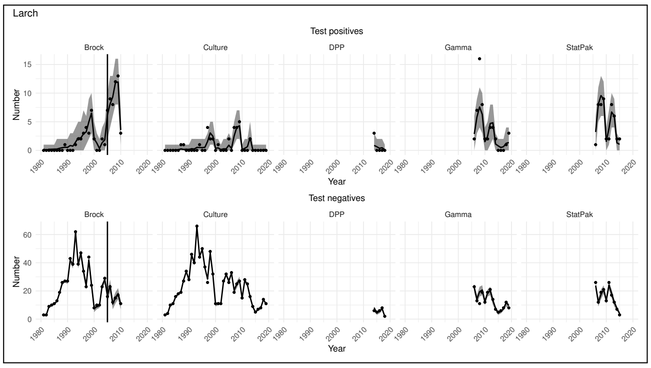
Larch
Test positives
Brock
Culture
DPP
Gamma
StatPak
Number
Year
Test negatives

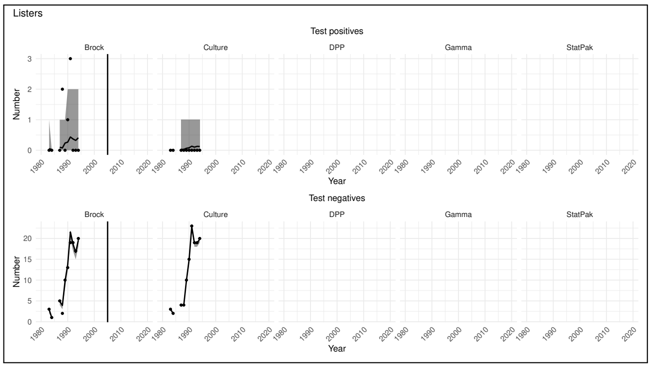
Listers
Test positives
Brock
Culture
DPP
Gamma
StatPak
Number
Year
Test negatives

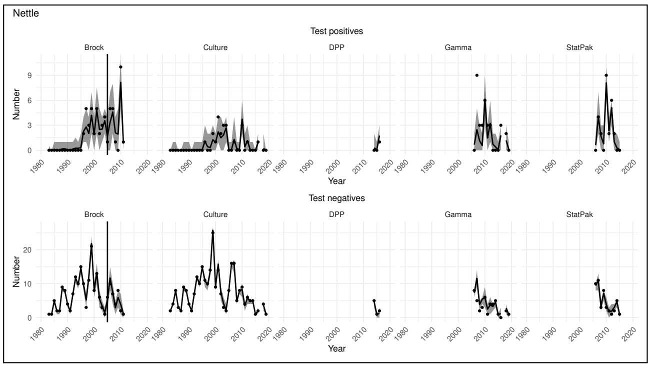
Nettle
Test positives
Brock
Culture
DPP
Gamma
StatPak
Number
Year
Test negatives

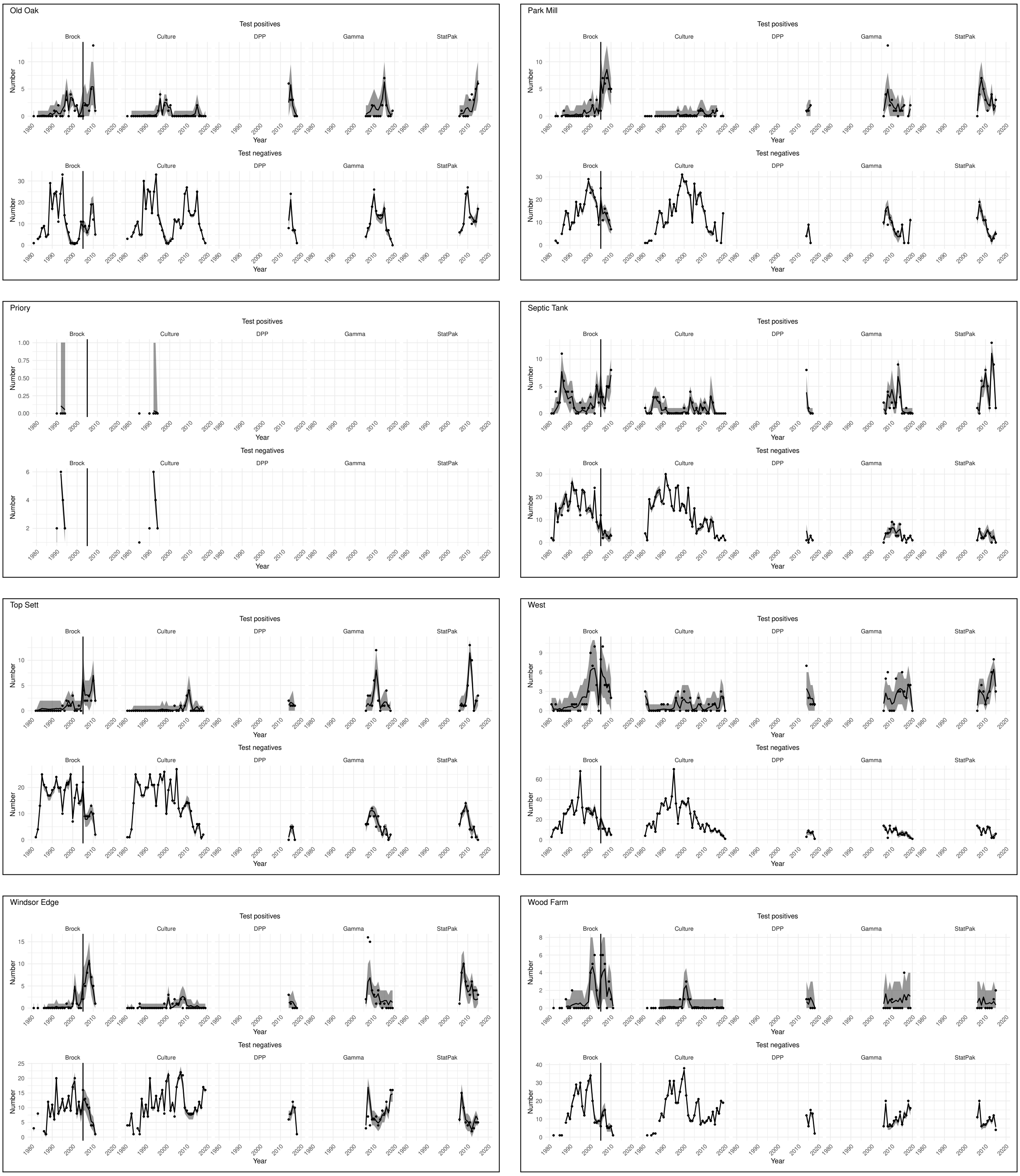
Old Oak
Park Mill
Priory
Septic Tank
Top Sett
West
Windsor Edge
Wood Farm
Test positives
Test negatives
Brock
Culture
DPP
Gamma
StatPak
Number
Year

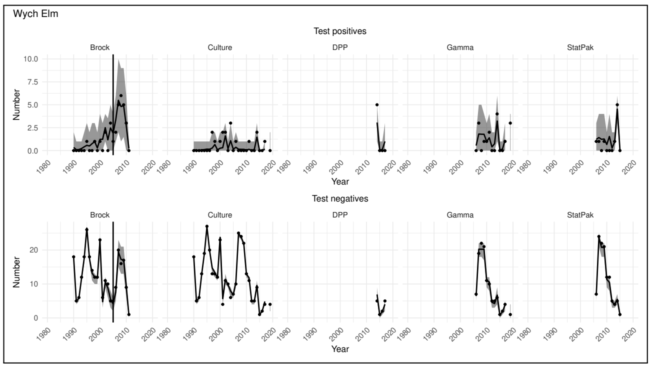
Wych Elm
Test positives
Brock
Culture
DPP
Gamma
StatPak
Number
Year
Test negatives

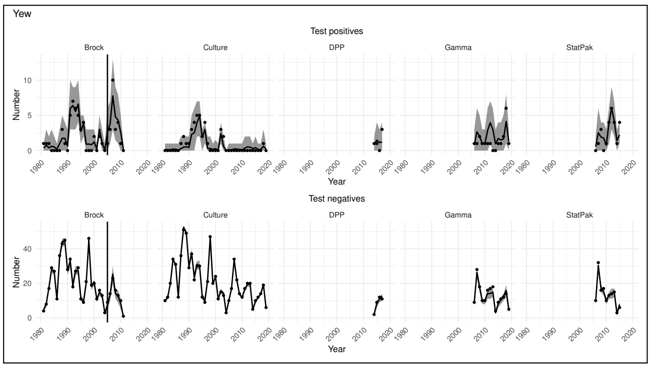
Yew
Test positives
Brock
Culture
DPP
Gamma
StatPak
Number
Year
Test negatives

#### A7.4 Results for model without Brock changepoint

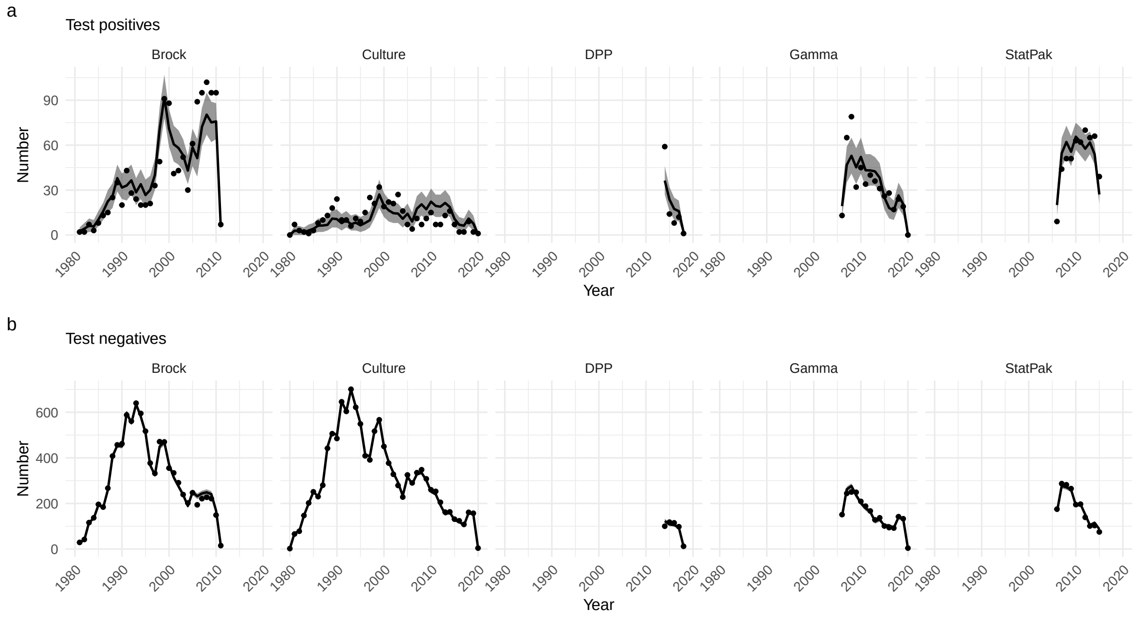

Figure A5: Posterior predictive distribution for the number of (A) test-positive and (B) test-negative results in each year against the observed data. Posterior means are indicated by black lines and 95% credible intervals by grey ribbons. Data are represented by black points.

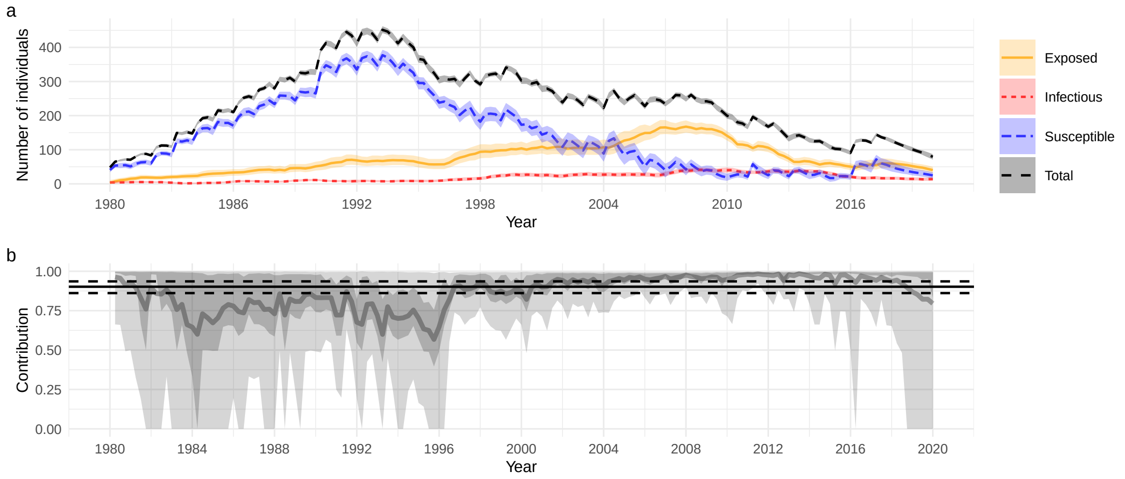

Figure A6: Posterior distribution (mean and 95% credible intervals) of the number of individuals in each infection state.

| Type | Parameter | Mean | 95% CI |
|---|---|---|---|
| Transmission | $\beta$ | 0.092 | (0.041, 0.2) |
| | $q$ | 0.64 | (0.19, 0.97) |
| | $\tau$ (years) | 7 | (5.9, 8.5) |
| Mortality | $a$ | 0.036 | (0.032, 0.039) |
| | $b$ | 0.033 | (0.029, 0.037) |
| | $c$ | 0.0016 | (0.000036, 0.0058) |
| Sensitivity ($E$) | Brock | 0.17 | (0.14, 0.19) |
| | Culture | 0.015 | (0.011, 0.02) |
| | DPP | 0.072 | (0.039, 0.11) |
| | Gamma | 0.11 | (0.096, 0.13) |
| | StatPak | 0.078 | (0.063, 0.095) |
| Sensitivity ($I$) | Brock | 0.88 | (0.84, 0.91) |
| | Culture | 0.32 | (0.29, 0.35) |
| | DPP | 0.63 | (0.53, 0.74) |
| | Gamma | 0.57 | (0.53, 0.62) |
| | StatPak | 0.93 | (0.89, 0.96) |
| Specificity | Brock | 0.99 | (0.99, 0.99) |
| | Culture | 1 | (0.99, 1) |
| | DPP | 0.95 | (0.88, 1) |
| | Gamma | 0.99 | (0.98, 1) |
| | StatPak | 0.97 | (0.92, 1) |
| Capture Probability | Winter | 0.18 | (0.17, 0.18) |
| | Spring | 0.36 | (0.35, 0.37) |
| | Summer | 0.46 | (0.45, 0.47) |
| | Autumn | 0.27 | (0.27, 0.28) |

Table A2: Posterior means and 95% credible intervals for key model parameters (to 2 significant figures).

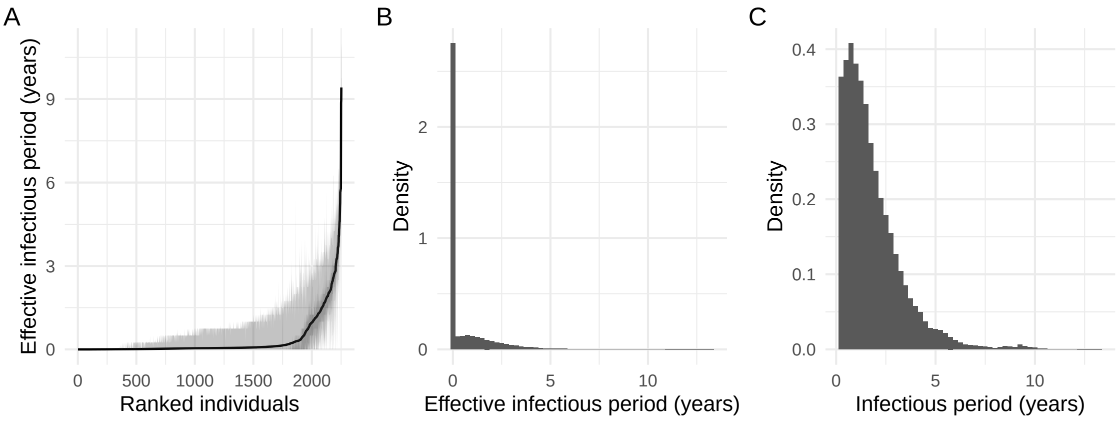

Figure A7: (A) Posterior means and credible intervals (50% and 95%) for the individual infectious period distribution for all individuals, ranked in increasing order of posterior means, (B) posterior distribution for the population-averaged infectious period distribution.

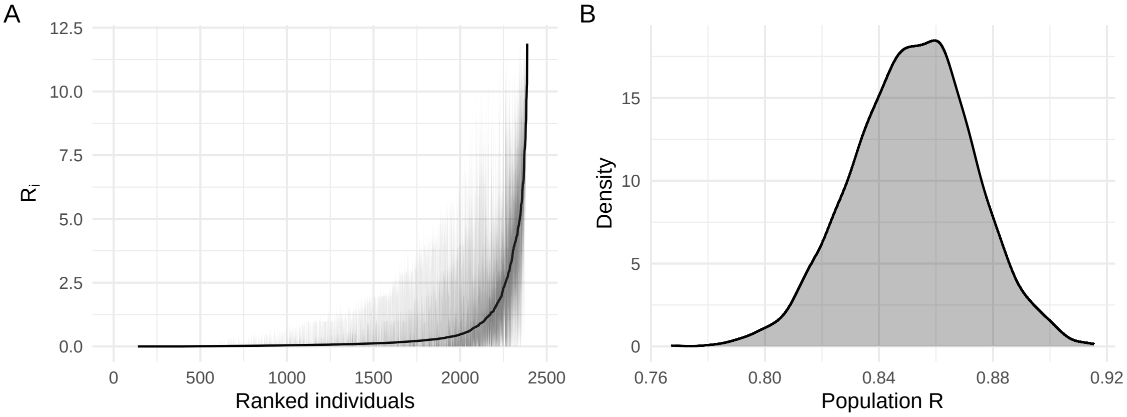

Figure A8: (A) Posterior means and credible intervals (50% and 95%) for the individual reproduction number for all individuals, ranked in increasing order of posterior means, (B) posterior distribution for the population reproduction number.

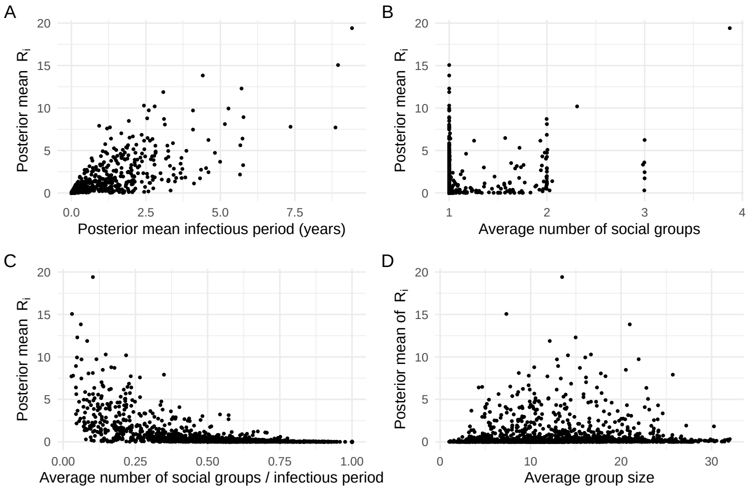

Figure A9: Posterior mean estimates of the individual reproduction numbers against (A) posterior mean duration of the infectious period, (B) average number of social groups an individual belonged to, (C) average number of social groups an individual belonged to divided by the length of the infectious period, and (D) the average group size for social groups that the individual belonged to.
